# Fine-tuning of outer membrane–peptidoglycan tethering by the redox-active lipoprotein LppB from Salmonella enterica

**DOI:** 10.1101/2025.11.13.688167

**Authors:** Elisa S. Pierre Despas, Seung-Hyun Cho, Bogdan I. Iorga, Jean-François Collet

## Abstract

The bacterial envelope is an essential compartment for survival, acting as a protective and permeability barrier. In diderm bacteria, it comprises an inner membrane, a thin peptidoglycan layer, and an outer membrane. A stable outer membrane-peptidoglycan connection is crucial for envelope integrity, preventing outer membrane detachment and enabling the generation of periplasmic turgor that counterbalances cytoplasmic pressure, thereby limiting susceptibility to antibiotics and environmental stressors. In *Escherichia coli*, the major lipoprotein Lpp (Braun’s lipoprotein) serves as the only covalent linker between the outer membrane and peptidoglycan, through its unique C-terminal lysine residue. Homologs of Lpp are widely conserved among Enterobacteriaceae, including *Salmonella enterica* serovar Typhimurium, which encodes LppA and LppB. While Lpp and LppA function similarly, the role of LppB remains unclear. Using *E. coli* as a surrogate model, we investigated the structural and functional properties of *Salmonella* LppB. Here, we demonstrate that LppB forms disulfide-linked dimers via its unique C-terminal cysteine and crosslinks to peptidoglycan through its terminal lysine, albeit with significantly lower efficiency than Lpp. Notably, LppB interacts with Lpp to form heterotrimers, thereby reducing overall Lpp-peptidoglycan crosslinking. These findings suggest that LppB modulates outer membrane-peptidoglycan tethering by fine-tuning the bacterial envelope architecture in response to environmental conditions. Given the established link between outer membrane-peptidoglycan uncoupling and outer membrane vesicles (OMV) formation, a key mechanism in bacterial pathogenicity, our results suggest a model in which LppB regulates outer membrane-peptidoglycan attachment, potentially influencing OMV production. This study highlights a modulating role for LppB in envelope remodeling, providing new insights into how *Salmonella* adapts its envelope structure in response to environmental cues to enhance fitness and virulence.

## INTRODUCTION

The cell envelope of Gram-negative bacteria is a complex structure essential for viability, providing the first line of defense against external threats. Its integrity must be preserved for bacteria to withstand diverse environmental stresses. The envelope comprises three distinct layers: an inner and an outer membrane that delimit a compartment known as the periplasm, in which lies the peptidoglycan ^1^. While the inner membrane is a classical phospholipid bilayer, the outer membrane is asymmetric, with phospholipids in the inner leaflet and lipopolysaccharides (LPS) in the outer leaflet. This asymmetry, together with the chemical properties of LPS, enables the outer membrane to function as an efficient permeability barrier^2^. In addition, the outer membrane contributes to the mechanical integrity of the cell envelope, functioning as a load-bearing structure ^3^. The peptidoglycan is a mesh-like polymer composed of glycan strands crosslinked by short peptides, which confers shape, mechanical strength, and osmotic resistance to the cell ^4^. The outer membrane is attached to the peptidoglycan, and this attachment has emerged as a key determinant of envelope stability and bacterial fitness ^5–7^. More recently, we demonstrated that proper attachment of the outer membrane to the peptidoglycan is required for resistance to hypoosmotic shock: this connection enables the build-up of periplasmic turgor, which counterbalances cytoplasmic pressure and protects the integrity of the inner membrane ^8^.

In *Escherichia coli*, outer membrane attachment to the peptidoglycan is primarily mediated by three proteins: Lpp, which forms a covalent bond to the peptidoglycan, and OmpA and Pal, which interact through non-covalent contacts ^5,9,10^. Lpp is a small α-helical lipoprotein that assembles into trimeric coiled-coil structures ^11–13^. It provides the only covalent attachment between the outer membrane and the peptidoglycan and plays a central role in maintaining envelope architecture ^14^. Remarkably, modifying the length of Lpp alters the width of the periplasm and changes the spacing between the two membranes ^15,16^. Lpp, the first protein identified as a lipoprotein ^5^, carries an N-terminal lipid moiety that anchors it in the inner leaflet of the outer membrane. After Sec-dependent translocation to the periplasm, immature Lpp undergoes three maturation steps: diacylation of the N-terminal cysteine, signal peptide cleavage, and addition of a third acyl chain to the same cysteine ^17,18^. Mature Lpp is then extracted from the inner membrane by the Lol machinery and delivered to the inner leaflet of the outer membrane ^19–21^. With more than one million copies per cell, Lpp is the most abundant protein in *E. coli* ^22^, and about one-third of this pool is covalently attached to the peptidoglycan ^6^. This attachment is catalyzed by the L,D-transpeptidases LdtA, LdtB, and LdtC, which link the ε-amino group of the C-terminal lysine of Lpp to the α-carboxyl group of the meso-diaminopimelic acid (mDAP) at position three of the peptidoglycan stem peptide ^23^.

Shortly after its discovery in *E. coli*, homologs of Lpp were identified in other closely related γ- proteobacterial families, including the *Enterobacteriaceae*, *Vibrionaceae*, and *Pseudomonadaceae* ^24^. Among these, the pathogenic species *Salmonella enterica* serovars Typhimurium and Typhi—two close relatives of *E. coli* that cause salmonellosis and typhoid fever, respectively—encode *lppA*, a gene whose product is nearly identical to *E. coli* Lpp, differing by only a single amino acid in the mature protein sequence (Fig. 1A). In *Salmonella*, LppA functions as the main outer membrane–peptidoglycan connector, mirroring the role of Lpp in *E. coli* ^16^. Adjacent to *lppA* lies a second homolog, *lppB* (Fig. 1B). LppB retains the conserved C-terminal lysine required for covalent peptidoglycan attachment ^25^, but diverges in the six residues preceding this lysine, notably featuring a cysteine at the penultimate position—an unusual trait given the high reactivity of cysteine in the oxidizing periplasm (Fig. 1A). *lppB* is expressed only at very low levels under standard laboratory conditions and during intracellular growth in macrophages ^26,27^, and its physiological significance remains uncertain. Nonetheless, previous studies have linked *lppB* to *Salmonella* virulence ^28,29^.

**Fig 1.**
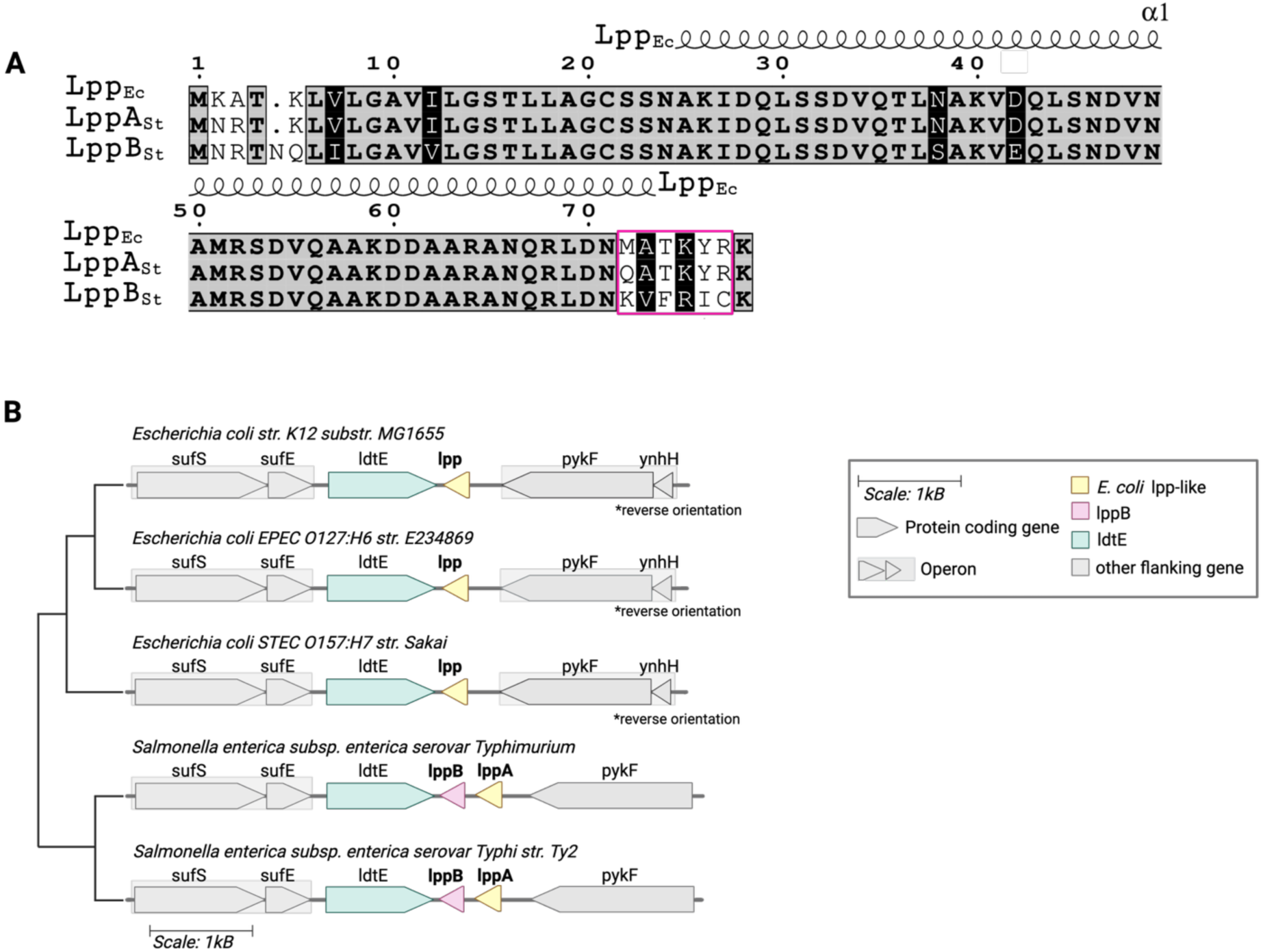
LppB shares a high degree of homology with Lpp and LppA but features a cysteine residue in the penultimate position of the C-terminal lysine. **A.** Amino acids sequence alignment of Lpp proteins of *E. coli* and *S.* Typhimurium LT2. The six residues preceding the C-terminal lysine (Lys58 in the matured peptide sequence of LppB starting from the tri-acylated cysteine, Cys21) are indicated within a pink box. Numbers above the first sequence line indicate amino acid positions relative to the start of each protein sequence. Conserved residues are shown as black letters on a grey background, and similar residues are shown as white letters in black boxes. The secondary structure of *E. coli* Lpp is depicted above the alignment. **B.** Schematic comparison of the genetic neighborhood of *lpp* in *E. coli*, *S.* Typhi, and *S.* Typhimurium. The *lpp* and *ldtE* genes face each other in both *E. coli* and *S. enterica* species and are positioned between the *suf* (iron-sulfur cluster) and *pykF* operons. LdtE catalyzes the formation of 3-3 crosslinks in the peptidoglycan ^48^. Phylogeny is midpoint rooted. Genes and intergenic regions are drawn to scale. Arrow tags represent genes, with arrow orientation indicating transcriptional direction. Colors depict homologous relationships. Illustration created with ESPript and Biorender.com.

In this study, we examined the ability of LppB to crosslink to the peptidoglycan, investigated the functional role of its penultimate cysteine, and evaluated its contribution to outer membrane tethering. We found that, like Lpp and LppA, LppB can be covalently attached to the peptidoglycan via its C-terminal lysine, albeit with lower efficiency. Notably, the cysteine immediately upstream of this lysine is susceptible to oxidation, leading to the formation of a homodimeric disulfide. We also showed that LppB assembles into heterotrimers with Lpp, and that even low expression levels of LppB reduce Lpp’s crosslinking to the peptidoglycan. Together, these findings indicate that LppB functions as a negative regulator of the Lpp– peptidoglycan connection in *S.* Typhimurium, fine-tuning outer membrane-peptidoglycan tethering in response to environmental conditions.

## RESULTS

### LppB covalently crosslinks to the peptidoglycan via its conserved C-terminal lysine

Given the high degree of conservation in envelope architecture and peptidoglycan composition between *E. coli* and *Salmonella*, we reasoned that *E. coli* provides a suitable model to investigate the intrinsic properties of LppB. Using a Δ*lpp* strain eliminates competition from endogenous Lpp, enabling direct comparison of LppB and Lpp under identical cellular conditions. Plasmid-based expression further allows control of protein levels and targeted mutagenesis, facilitating a functional dissection of LppB crosslinking to the peptidoglycan.

We first tested whether the C-terminal lysine of LppB can be covalently linked to the meso-diaminopimelic acid (mDAP) residue at position three of the peptidoglycan stem peptide. To assess peptidoglycan crosslinking, lysates of lysozyme-treated cells were analyzed by SDS-PAGE and Western blot using affinity-purified polyclonal anti-Lpp antibodies (95% identity with the corresponding LppB epitope) (Fig. 2). As described previously ^30^, lysozyme digests the glycan chains of the peptidoglycan, fragmenting the cell wall and releasing covalently attached proteins. This treatment is essential to recover peptidoglycan-bound forms of Lpp or LppB, which are otherwise too large to migrate into the gel and thus remain undetectable. The appearance of high-molecular-weight bands following lysozyme treatment therefore provides a readout of covalent crosslinking. As expected, Lpp produced high-molecular-weight bands (Fig. 2, lanes 6–7). Similar bands were observed for LppB (Fig. 2, lane 9) but were abolished when its C-terminal lysine was mutated to arginine (K58R) (Fig. 2, lane 10). Thus, LppB, like Lpp, is covalently crosslinked to the peptidoglycan through its terminal lysine. Interestingly, prominent bands of ∼15 kDa were consistently detected in cells expressing LppB (Fig. 2, lanes 4–5; 9–10), but not in those expressing Lpp, suggesting that LppB forms homodimers under physiological conditions (see specific section below and Fig. 4).

**Figure 2.**
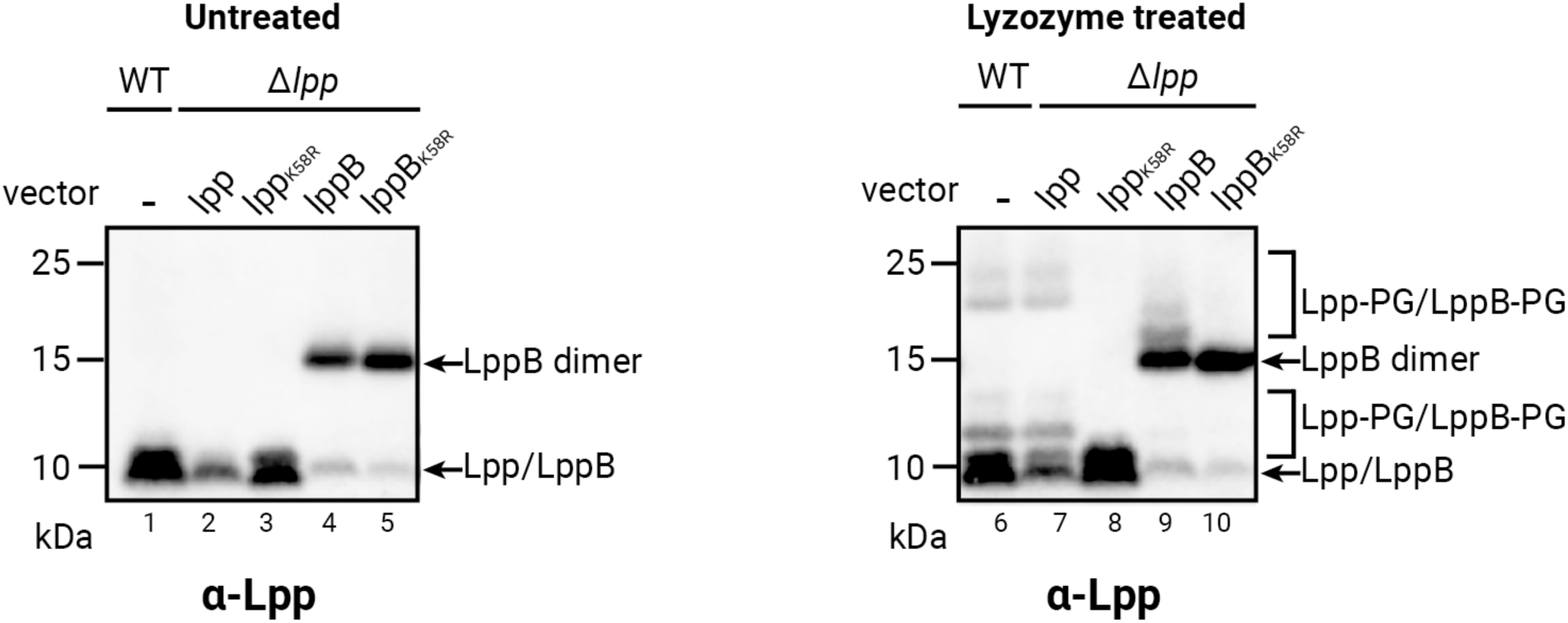
LppB crosslinks to the peptidoglycan via K58. LppB crosslinking to peptidoglycan (PG). Lpp and LppB variants were expressed from pSC284 and pSC283 derivatives, respectively, in the Δ*lpp* strain (lanes 2-5; 7-10). The left panel depicts untreated cell lysate (lanes 1-5) while the right panel depicts cell lysates treated with 25 μg/μL of lysozyme (lanes 6-10). Bands corresponding to the monomeric form of Lpp and LppB are observed at ∼7.5 kDa. Bands corresponding to the dimeric form of LppB are observed at ∼ 15 kDa. Bands of higher molecular weight corresponding to Lpp or LppB bound to various muropeptides are observed except in strains expressing the K58R variant of Lpp or LppB (lanes 8 and 10 respectively). Cell lysates were TCA-precipitated and Western blot analysis was performed using anti-Lpp antibody. Representative images from experiments conducted in biological triplicates are shown.

Since covalent attachment of Lpp to the peptidoglycan contributes to outer membrane stability, we next considered how conditions relevant to infection might influence this process. Within macrophage phagosomes, *Salmonella* encounters an acidic environment (pH 4.0–5.0), which it not only tolerates but exploits to activate virulence programs and promote intracellular survival ^31,32^. Based on this, we hypothesized that LppB expression and crosslinking activity could be modulated by pH. Indeed, both Lpp and LppB showed increased expression at acidic pH, regardless of whether they were expressed from the chromosome or a plasmid (Fig. S1). Consistently, LppB complemented the SDS sensitivity of the Δ*lpp* strain more effectively at pH 5.5 than at neutral pH (Fig. S2). These results indicate that acidic conditions enhance LppB expression and functional contribution to envelope integrity. To reflect more accurately the intraphagosomal conditions experienced by *Salmonella* during infection, all subsequent experiments were therefore conducted at acidic pH.

### LdtB is essential for efficient peptidoglycan crosslinking of LppB

We then asked whether the three L,D-transpeptidases—LdtA, LdtB, and LdtC—dedicated to Lpp crosslinking in *E. coli* can also catalyze the attachment of LppB to the peptidoglycan. These enzymes are also present in *Salmonella*, where they share substantial (78-93%) sequence identity with their *E. coli* counterparts ^33^ (Fig. S3). To address this question, we analyzed strains carrying individual deletions of each *ldt* gene and examined their crosslinking profiles. LppB crosslinking was lost in the absence of LdtB, whereas deletion of *ldtA* or *ldtC* had no detectable effect (Fig. 3A, lanes 4–6). Because LdtA and LdtC are expressed at much lower levels than LdtB (10- and 100-fold less abundant, respectively ^22^), we tested whether their overexpression could promote LppB attachment. To this end, we expressed *ldtA* or *ldtC* from a lac promoter– based plasmid in a Δ*ldtB* background and analyzed crosslinking. Overexpression of LdtA partially restored high–molecular-weight LppB species (Fig. 3B, lanes 11 and 18), whereas overexpression of LdtC showed little or no activity (Fig. 3B, lanes 13 and 20). As expected, efficient crosslinking was restored when LdtB was provided in *trans* (Fig. 3B, lanes 12 and 19). Taken together, these results indicate that LdtB is the dominant enzyme responsible for LppB crosslinking, while LdtA can contribute weakly under overexpression conditions and LdtC appears inactive.

**Figure 3.**
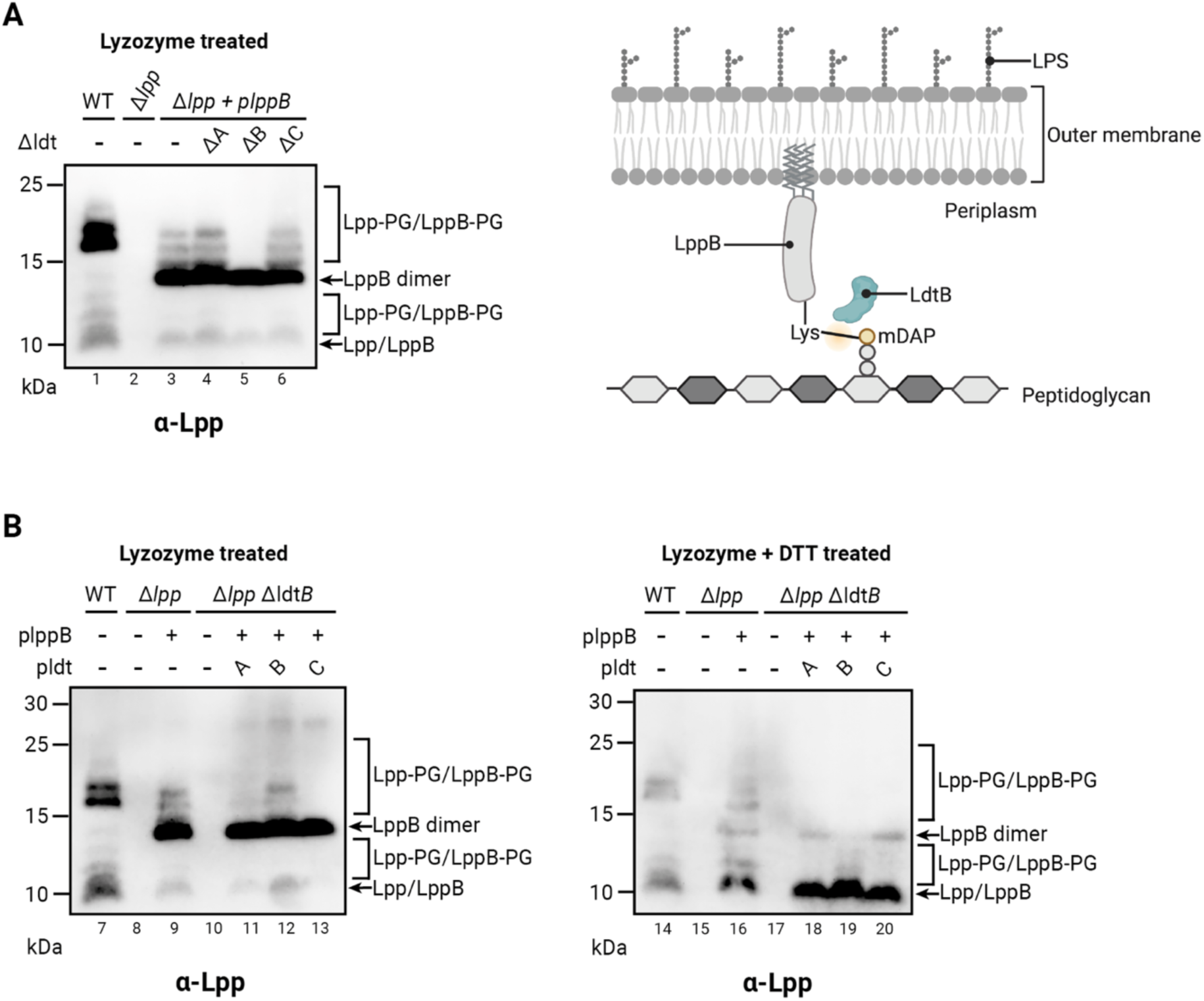
LdtB is the primary L,D-transpeptidase for efficient peptidoglycan crosslinking of LppB. LppB crosslinking to peptidoglycan (PG) in the presence or absence of the L,D-transpeptidases. **A**. LdtB is required for efficient LppB crosslinking. LppB was expressed from pSC283 (pLppB) in the Δ*lpp* mutant or in double mutants *Δlpp/ΔldtA*, *Δlpp/ΔldtB*, *Δlpp/ΔldtC* strains (lanes 3-6). Deletion of *ldtB* (lane 5) caused the strongest reduction of LppB-peptidoglycan crosslinking, whereas loss of *ldtA* or *ldtC* had only minor effects. Wild-type and *Δlpp* controls are shown in lanes 1 and 2, respectively. A schematic (right) illustrates the LdtB-mediated crosslink: the C-terminal lysine residue of LppB (K58) is covalently attached to the diaminopimelic acid (mDAP) moiety of the peptidoglycan. **B.** Functional complementation of LppB crosslinking in the absence of LdtB. LdtA (pEP25; lanes 11 and 18), LdtB (pAA186; lanes 12 and 19), and LdtC (pEP26; lanes 13 and 20) were ectopically expressed alongside LppB (pLppB) in the *Δlpp/ΔldtB* strain (lanes 10 and 17). Wild-type lanes (7 and 14), *Δlpp* lanes (8 and 15) and *Δlpp* complemented with pLppB lanes (9 and 16) served as controls. LdtB restoration (lanes 12 and 19) fully rescued crosslinking, while LdtA provided partial activity (lanes 11 and 18) and LdtC had minimal effect (lanes 13 and 20), confirming LdtB as the predominant transpeptidase. For both figures, sample were treated with 25 μg/μL of lysozyme, cell lysates were TCA-precipitated and treated with 30 mM DTT for reduction when indicated. Western blot analysis was performed using anti-Lpp antibody. Representative images from experiments conducted in biological triplicates are shown. Schematic illustration created with Biorender.com.

### LppB oxidation leads to homodimer formation via its cysteine residue

The C-terminal region of LppB differs markedly from that of Lpp and LppA. Notably, LppB carries a cysteine residue immediately upstream of the terminal lysine required for peptidoglycan crosslinking (Fig. 1B). Cysteine residues are highly reactive and can undergo oxidation to sulfenic acid, which may lead to disulfide bond formation or to irreversible modifications such as sulfinic or sulfonic acids ^34^. This is particularly relevant in the oxidizing ^32^ environment of the periplasm, where single cysteines are prone to oxidation ^35^. The presence of this cysteine in close proximity to the crosslinking site suggested that disulfide bond formation could interfere with peptidoglycan attachment. To test this, *lppB* was expressed in Δ*lpp* cells; after acid quenching and alkylation to prevent post-lysis oxidation ^36^, Western blot analysis revealed that LppB migrated predominantly as a ∼15 kDa species under non-reducing conditions (Fig. 4A, lane 5; Fig. 4B, lane 14), consistent with a disulfide-linked homodimer. This band disappeared upon addition of DTT (Fig. 4A, lane 10; Fig. 4B, lane 20), confirming its disulfide-dependent nature. In contrast, Lpp showed no mobility shift. Furthermore, mutation of the cysteine to arginine (LppB_C57R_) abolished the mobility shift, and the mutant migrated like Lpp (Fig. 4B). These results demonstrate that LppB undergoes oxidation in the periplasm and forms a homodimeric disulfide via Cys57. Interestingly, the intensity of LppB bands was consistently lower than that of Lpp, suggesting reduced crosslinking efficiency. Whether this reduced efficiency reflects differences in the C-terminal region of LppB or results from expression in *E. coli* rather than in its native *Salmonella* context remains unclear. Because the disulfide lies immediately adjacent to Lys58, we hypothesized that its formation could impair peptidoglycan crosslinking. Substitution of the cysteine with arginine (LppB_C57R_), mimicking the naturally occurring residue in Lpp and LppA, however, did not restore efficient crosslinking (Fig. 4B, compare lanes 18, 20, 22). The reduced expression of LppB_C57R_ complicates the interpretation, but the results indicate that removal of the disulfide bond alone does not restore efficient crosslinking to the peptidoglycan, suggesting that additional features of the LppB C-terminal region underlie its reduced efficiency.

**Figure 4.**
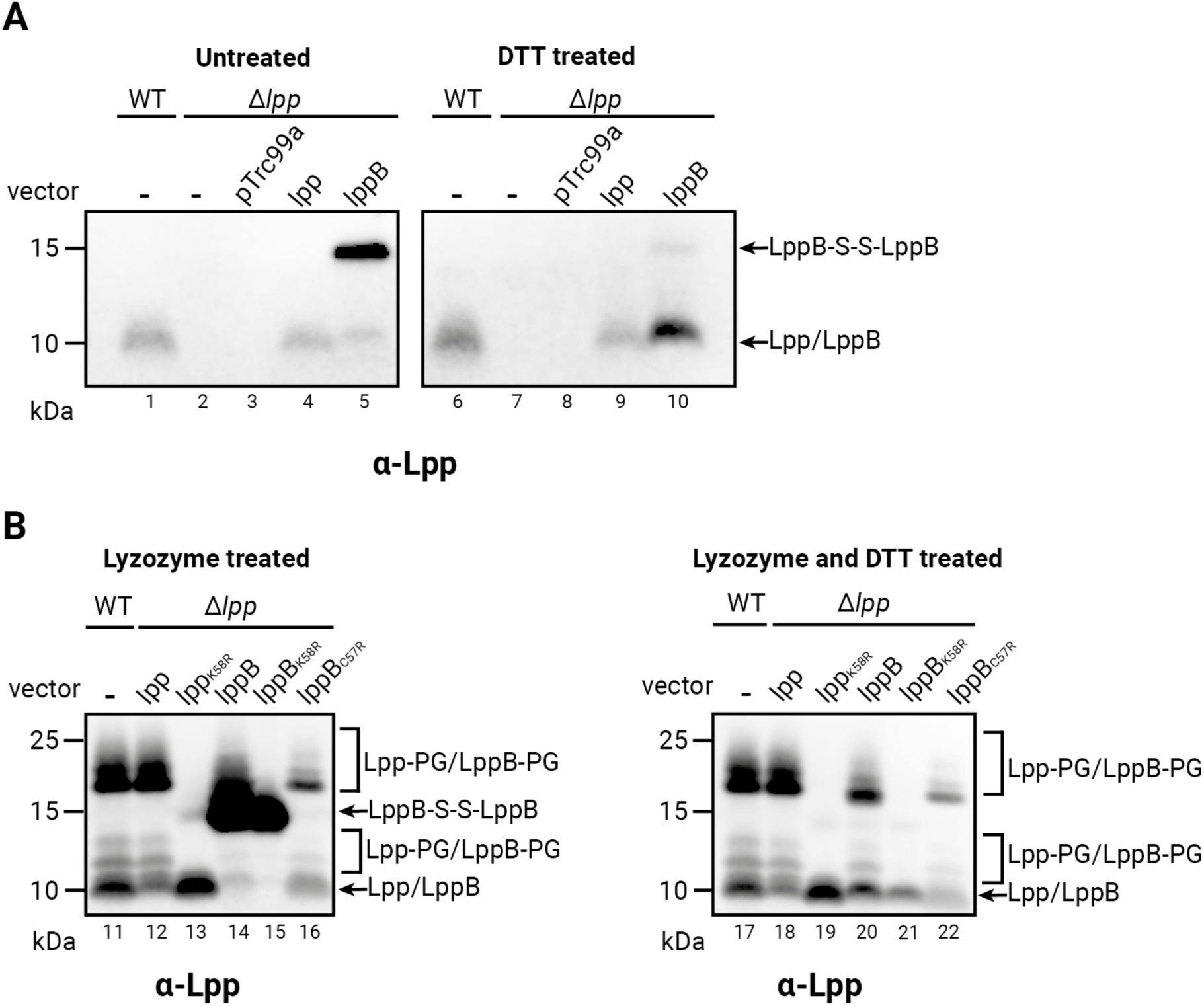
LppB predominantly forms disulfide-linked homodimers *in vivo*. LppB mostly exists in an oxidized state *in vivo*, forming mixed disulfide homodimers. Lpp and LppB variants were expressed from pSC284 and pSC283 derivatives, respectively, in the Δ*lpp* strain (lanes 4-5; 9-10; 12-16; 18-22). The wild type is shown in lanes 1, 6, 11 and 17 as control. **A.** Oxidized state of LppB *in vivo*. The left panel depicts untreated cell lysate while the right panel depicts cell lysate treated with 30 mM DTT to reduce disulfide bonds. Monomeric forms of Lpp and LppB migrate at ∼ 7.5 kDa, whereas bands corresponding to the dimeric form of LppB are observed at ∼ 15 kDa. DTT treatment reduced homodimer abundance with a corresponding increase of monomer signal (compare lane 5 and 10), confirming disulfide bond-mediated dimerization. **B.** Cysteine-dependent homodimerization of LppB. The left panel depicts cell lysates treated with 25 μg/μL of lysozyme (lanes 11-16). The right panel depicts samples additionally treated with 30 mM DTT to reduce disulfide bonds (lanes 17-22). High–molecular weight bands corresponding to Lpp bound to peptidoglycan fragments were observed, except in lanes carrying the lysine substitution mutants (K58R; lanes 13,15,19,21). Substitution of the cysteine (C57R) abolished LppB homodimeric species (LppB-S-S-LppB; compare lane 14 and 16). DTT treatment similarly reduced the LppB homodimer signal and increased monomer abundance (compare lane 14 and 20). Western blot analysis was performed using anti-Lpp antibody, and representative images from experiments conducted in biological triplicates are shown.

### Lpp and LppB form heterotrimeric complexes linked by disulfide bonds

The experiments described above were performed in the absence of Lpp. To determine whether Lpp influences LppB dimerization and peptidoglycan attachment, we expressed LppB in wild-type cells at both high and low levels. In all cases, LppB formed a stable disulfide-linked homodimer, identical to that observed in the Δ*lpp* strain (Fig. 4A, Fig. 5A, Fig. S4). Thus, LppB homodimerization occurs independently of expression level or the presence of Lpp. Given the high similarity between LppA and LppB (Fig. 1B) and the trimeric nature of Lpp, we then hypothesized that LppA and LppB might assemble into heterotrimers in *Salmonella*. To test this, we used *E. coli* Lpp as a proxy for LppA and employed two chemical crosslinkers, DSP [dithiobis(succinimidyl propionate)] and BS3 [bis(sulfosuccinimidyl) suberate], to stabilize potential Lpp–LppB interactions. Both reagents react with primary amines, but DSP contains an internal disulfide bond that is cleavable under reducing conditions, whereas BS3 generates irreversible crosslinks. Both crosslinkers efficiently produced crosslinked oligomeric species for Lpp and LppB (Fig. 5). For Lpp alone, the predominant crosslinked species was a dimer (Fig. 5B, lane 3; Fig. 5C, lane 7), consistent with earlier reports that amine-reactive crosslinkers preferentially capture Lpp dimers rather than Lpp trimers ^37,38^. For LppB alone, multiple disulfide bonds between homotrimers led mainly to non-crosslinked mixed disulfide species of LppB (LppB–S–S–LppB; Fig. 5B, lane 6; Fig. 5C, lane 10, Fig. 5G). Interestingly, after reduction of BS3-treated samples, we detected crosslinked LppB–LppB dimers (Fig. 5F, lane 20; Fig. 5G). In contrast, no such species were observed after DSP treatment (Fig. 5E, lane 16). Because DSP crosslinks are lost upon reduction, whereas BS3 crosslinks remain intact, this pattern suggests that the BS3-resistant bands correspond to LppB dimers that had been both disulfide-bonded and chemically crosslinked within higher-order LppB oligomers. When Lpp and LppB were co-expressed, two dimer-sized bands were detected (Fig. 5B, lanes 4–5; Fig. 5C, lanes 8–9), consistent with Lpp–Lpp and disulfide-linked LppB–LppB species (Fig. 5G). Because LppB predominantly forms disulfide-bonded homodimer, crosslinked Lpp-LppB heterodimer cannot be detected as a dimer-sized band. We therefore reasoned that if heterodimers formed, they might be stabilized as higher-order complexes such as Lpp–LppB–S–S–LppB–Lpp. Indeed, a band of the expected size was observed only in samples co-expressing Lpp and LppB (blue asterisks; Fig. 5B, lanes 4–5; Fig. 5C, lanes 8–9, Fig. 5G). Upon DTT reduction, the tetrameric bands observed in co-expressed Lpp/LppB samples disappeared (compare Fig. 5B, lanes 4–5 with Fig. 5E, lanes 14–15; Fig. 5C, lanes 8–9 with Fig. 5F, lanes 18–19), and strong dimeric species reappeared in BS3-treated samples (Lpp–X–LppB; brown asterisks; Fig. 5F, lanes 18– 19). To further characterize these dimers, we treated the reduced samples with the alkylating reagent AMS [4-acetamido-4`′`-maleimidylstilbene-2,2`′`-disulfonic acid], which covalently modifies free thiols by adding ∼500 Da. As expected, the crosslinked Lpp dimer did not shift upon AMS treatment, consistent with the absence of cysteine residues. In contrast, the LppB dimer displayed a mobility shift, indicating AMS modification of its cysteine. Importantly, the presumptive Lpp–LppB dimer also shifted after AMS treatment, confirming that it contains a cysteine residue contributed by LppB (Fig. S5). Altogether, these results support the formation of heteromeric complexes between Lpp and LppB *in vivo*, with heterotrimers connected by mixed disulfide bonds involving the cysteine of LppB. Given that LppB is likely expressed at lower levels than LppA in *Salmonella*, we propose that the predominant native heterotrimer consists of two LppA subunits and one LppB subunit, arranged as [LppA₂LppB]–S–S– [LppBLppA₂] (Fig. 5H).

**Figure 5.**
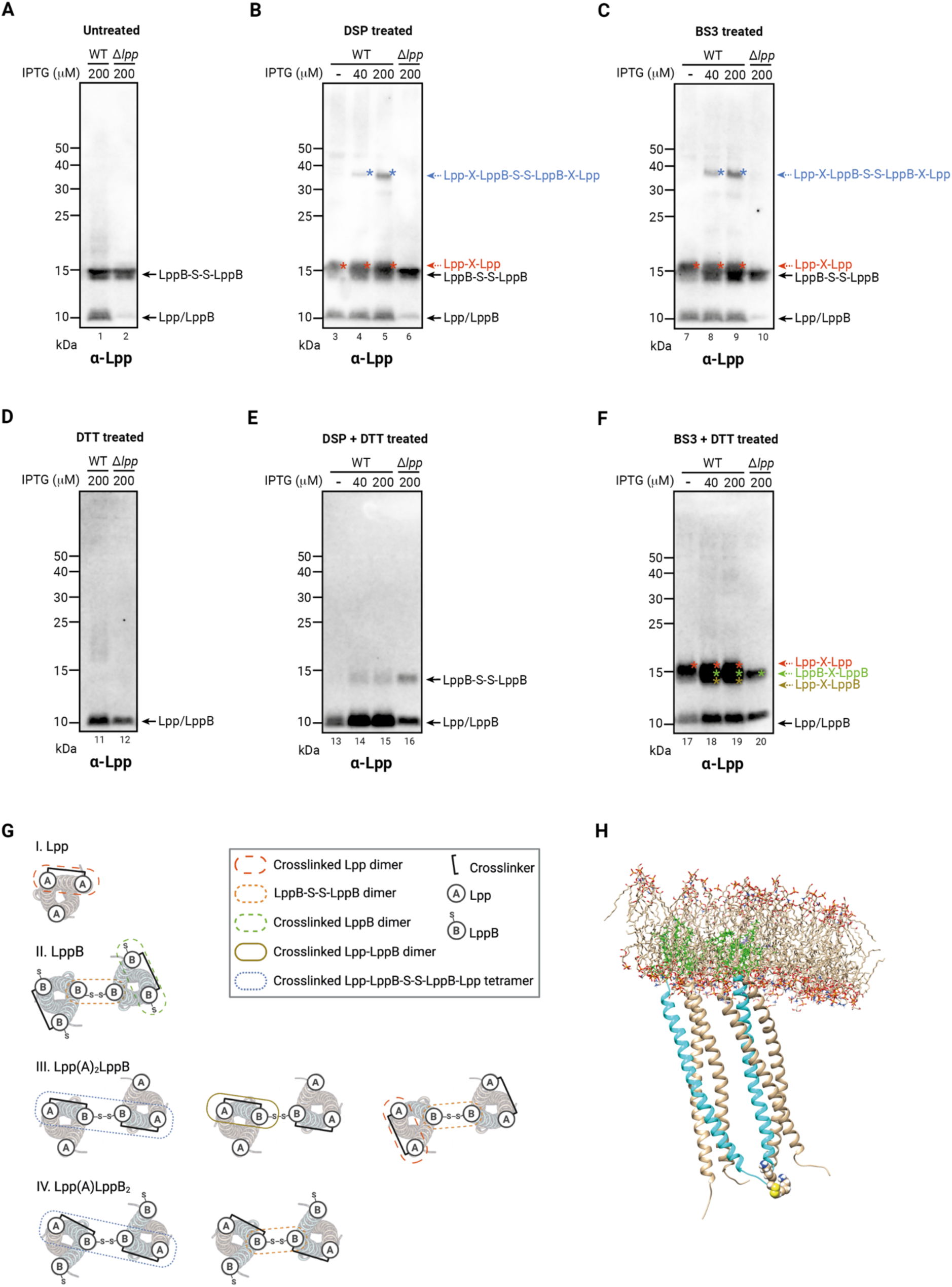
LppB drives disulfide-linked dimerization of Lpp/LppB heterotrimers *in vivo*. LppB was expressed from pSC283 (pLppB) in the wild-type and *Δlpp* strains using varying isopropyl ß-D-1-thiogalactopyranoside (IPTG) concentrations for induction. **A.** LppB forms a homodimeric mixed disulfide in the presence or absence of Lpp **B. -C.** Chemical crosslinking with Dithiobis-succinimidyl-propionate (DSP) or with Bis-sulfosuccinimidyl-suberate (BS3) reveals higher order oligomers. Distinct bands corresponding to Lpp homodimers and to Lpp-LppB heterotetramers (Lpp-LppB-S-S-LppB-Lpp) are observed. **D.** Reducing conditions disrupt disulfide-linked oligomers. Treatment of samples from A. with 30 mM DTT decreased homodimer abundance and increased monomer signal. **E.** Reducing conditions on DSP-crosslinked complexes disrupts both disulfide-linked and chemically crosslinked homodimers and heterotetramers. Treatment of samples from B. with 30 mM DTT results in monomer enrichment and decrease of higher order oligomers. **F.** Reducing conditions on BS3-crosslinked complexes only disrupts disulfide-linked homodimers and heterotetramers. Treatment of samples from C. with 30 mM DTT results in monomers and dimers (LppB homodimers and Lpp-LppB heterodimers) enrichment and disulfide-linked heterotetramers (Lpp-LppB-S-S-LppB-Lpp) and homodimers (LppB-S-S-LppB) decrease. **G**. Schematic representation of oligomeric states of Lpp (represented as A) and LppB (represented as B) observed in panels A-F. Dimeric and tetrameric crosslinked species are depicted. **H.** Structural model of an Lpp-LppB heterohexamer. Two LppA-LppB mixed disulfide trimers [LppA_2_LppB] are linked via a disulfide bond to form a hexameric complex [LppA_2_LppB]-S-S-[LppBLppA_2_] embedded in the OM. The two LppB helices appear in cyan, with disulfide-bonded Cys57s and Lys58s displayed as spheric atoms (sulfur atoms in yellow; nitrogen atoms in the amino group in blue). N-terminal acyl chains (green) are embedded in the OM (shown as stick).

### LppB negatively modulates Lpp crosslinking to PG

Our results above showed that LppB exhibits reduced crosslinking to the peptidoglycan and can assemble into heterotrimers with Lpp. Based on these observations, we hypothesized that Lpp–LppB heterotrimers may also display reduced crosslinking efficiency, reflecting the weaker attachment of LppB to the peptidoglycan (Figure 4). Since LppB is likely expressed at lower levels than LppA in *S.* Typhimurium, the formation of such heterotrimers could nevertheless be frequent. Simple calculations indicate that if LppB represents only 10% of the total pool of Lpps, heterotrimers would still account for more than 27% of the total Lpp population (Supplementary Data). We therefore proposed that LppB might act as a negative modulator of LppA crosslinking. To test this, we expressed LppB at gradually increasing levels and monitored Lpp crosslinking to the peptidoglycan following lysozyme treatment. Increasing LppB expression led to a proportional decrease in the intensity of Lpp–peptidoglycan crosslinked bands (Fig. 6A), consistent with a dominant-negative effect of LppB. This effect was already detectable at low isopropyl ß-D-1-thiogalactopyranoside (IPTG) concentrations (10 μM), when LppB levels were much lower than those of Lpp (Fig. 6A, lane 2). Moreover, it persisted after DTT treatment, indicating that it is independent of the disulfide bond in LppB (Fig. 6B). To further validate this effect, we performed experiments at a fixed IPTG concentration (40 μM), comparing samples with and without induction. Again, a clear decrease in Lpp–peptidoglycan crosslinking was observed in the presence of LppB (Fig. 6C, Fig. S6). Importantly, no decrease was detected when Lpp was overexpressed from a plasmid in wild-type cells under comparable conditions, confirming that the effect is specific to LppB. Together, these findings indicate that even at low expression levels, LppB can negatively modulate Lpp crosslinking to the peptidoglycan *in vivo*.

**Figure 6.**
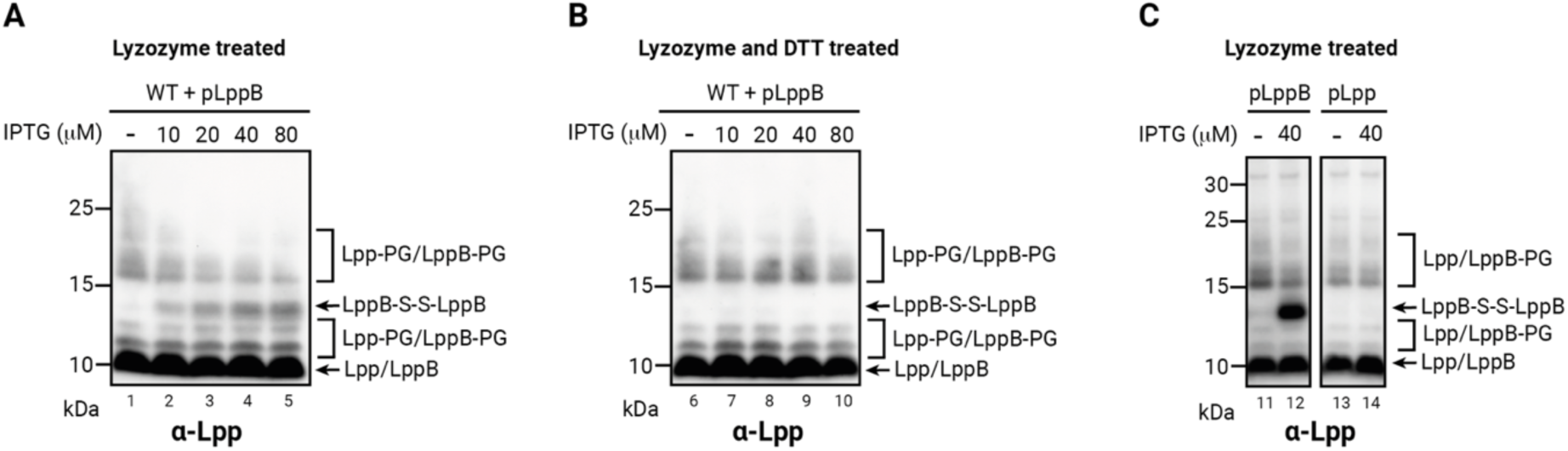
LppB negatively modulates Lpp-PG crosslinking. LppB was expressed from pSC283 (pLppB) in wild-type cells using the indicated isopropyl ß-D-1-thiogalactopyranoside (IPTG) concentrations for induction. Cells were harvested and treated with lysozyme for peptidoglycan digestion. **A.** Dose-dependent inhibition of Lpp-PG crosslinking by LdtB. Progressive induction of LppB reduced the abundance of Lpp–PG crosslinked species. **B.** Effect of reducing conditions on Lpp-PG crosslinking. Samples from panel A were additionally treated with 30 mM DTT to reduce disulfide bonds. The inhibitory effect of LppB on Lpp–PG crosslinking persisted after reduction, demonstrating that this modulating activity is independent of the disulfide bond in LppB. **C.** Comparison of LppB and Lpp expression effect on Lpp-PG crosslinking. LppB expression decreases Lpp–PG crosslinking, whereas Lpp expression does not. Left panel: LppB expressed from pSC283 (pLppB) at 0 or 40 μM IPTG. Right panel: Lpp expressed from pSC284 (pLpp) under the same conditions. Cells were harvested and treated with lysozyme for PG digestion. Western blot analysis was performed using anti-Lpp antibody. Representative images from experiments conducted in biological triplicates are shown. (see Fig. S6).

## DISCUSSION

In this study, using *E. coli* as a surrogate, we examined how *Salmonella* exploits the unique structural features of LppB—most notably its penultimate cysteine residue and capacity to form disulfide bonds—to modulate envelope integrity. We found that LppB, like Lpp and LppA, can crosslink to the peptidoglycan via its C-terminal lysine, but with markedly lower efficiency. In addition, LppB assembles into heterotrimers with Lpp, further influencing outer membrane– peptidoglycan connectivity by limiting lysine accessibility within hexameric complexes. Even when expressed at low levels, LppB reduced the extent of Lpp-mediated crosslinking, acting as a dampening factor that fine-tunes the strength of the outer membrane–peptidoglycan connection.

Our results support the view that the distinctive properties of LppB arise from two main features: (1) the presence of a penultimate cysteine residue, and (2) sequence variation in the six amino acids preceding the terminal lysine. The disulfide bond formed by Cys57 may hinder the access of L,D-transpeptidases to Lys58, while changes in the surrounding residues could reduce enzymatic efficiency, given that the corresponding stretch in Lpp/LppA is likely optimized for crosslinking. Despite this restriction, Lys58 remains conserved in LppB. Molecular dynamics suggest that, in heterotrimers, the disulfide bond restricts the mobility of LppB while stabilizing hexamers in a vertical orientation against the peptidoglycan (Supplementary Movie). This conformation may allow some crosslinking, albeit less efficient than in Lpp/LppA homotrimers, thereby explaining the conservation of Lys58.

The importance of cysteine residues in Lpp-like proteins is underscored by YqhH, an Lpp paralog in *E. coli*. YqhH contains two C-terminal cysteines, one of which aligns with the cysteine of LppB, but lacks a terminal lysine and therefore cannot crosslink to the peptidoglycan (Fig. S7). While its function remains unclear, YqhH may form complexes with Lpp and modulate crosslinking in a manner analogous to LppB, though via a distinct mechanism.

The impact of heterotrimer formation on envelope connectivity could be substantial. Calculations indicate that if LppB accounts for only 10% of total Lpps, heterotrimers would represent more than 27% of the population (Supplementary Data). Because heterotrimers are less efficiently crosslinked than homotrimers, this shift would weaken outer membrane– peptidoglycan attachment. Clustering of Lpp/LppB heterotrimers could create local zones of disconnection, altering periplasmic turgor balance and promoting outer membrane vesicle (OMV) formation ^8,39^. Since OMV production contributes to virulence and interspecies interactions, our findings suggest that LppB may play a role in envelope remodeling processes that support *Salmonella* fitness and pathogenicity.

Finally, our results show that LppB is predominantly oxidized *in vivo*. We did not determine whether disulfide bonds are introduced before or after peptidoglycan crosslinking, but the timing of this event could be critical for Ldt accessibility. The periplasmic thiol oxidase DsbA, which catalyzes disulfide bond formation ^40^, is a likely candidate to drive this process. Future work should investigate whether DsbA-dependent oxidation governs LppB function and thereby contributes to the regulation of outer membrane–peptidoglycan connectivity.

## MATERIALS AND METHODS

### Bacterial Strains and Plasmids

The bacterial strains and primers used in this study are listed in Table S1 and Table S3, respectively. The parental *E. coli* strain DH300 (a derivative of *E. coli* MG1655 carrying a chromosomal *rprAp::lacZ* fusion at the λ attachment site to monitor Rcs activation) was used as wild type throughout the study ^41^. Deletion mutants were generated by transferring the corresponding alleles from the Keio collection (kan^R^) ^42^ into DH300 by P1 phage transduction and verified by PCR.

The plasmids used in this study are listed in Table S2. To generate pSC283 and pSC284, *lppB* and *lpp* were PCR-amplified using NcoI_stLppB_Fw / XbaI_stLppB_stop_Rv and NcoI_ecLpp_Fw / XbaI_ecLpp_stop_Rv as primer pairs and chromosomal *S.* Typhimurium LT2 and *E. coli* DH300 DNA as templates, respectively. The PCR product was subsequently inserted into pTrc99a restricted with NcoI and XbaI. To introduce the *lppB* and *lpp* point mutation(s) at selected positions, site-directed mutagenesis was performed on pSC283 and pSC284 using the primers described in Table S3. To generate pEP25 and pEP26, *ldtA* and *ldtC* were PCR-amplified using BamHI_ecLdtA_Fw / PstI_ecLdtA_stop_Rv and KpnI_ecLdtC_Fw / PstI_ecLdtC_stop_Rv as primer pairs and chromosomal *E. coli* MG1655 DNA serving as a template, respectively. The PCR products were subsequently inserted into the low-copy vector pAM238 after enzymatic restriction by BamHI and PstI for *ldtA* and KpnI and PstI for *ldtC* ^43^.

### Bacterial culture and Protein expression

Bacterial strains used in this study are listed in Table S1. Bacterial strains were cultured in Miller’s Lysogeny Broth (LB Miller – NaCl 10g/L, Tryptone 10 g/L, Yeast Extract 10 g/L) at 37°C unless stated otherwise. When appropriate, antibiotics were used at the following concentrations: ampicillin 200 μg/mL; spectinomycin 100 μg/mL; kanamycin 50 μg/mL. Protein expression was achieved by inoculating fresh LB (pH 7.0 or pH 5.5) with an overnight culture, diluted at either 1:250 or 1:1000. Lpp, LppB, and their mutant proteins were expressed from pLpp (pSC284), pLppB (pSC283), and their derivatives by adding indicated concentration of isopropyl ß-D-1-thiogalactopyranoside (IPTG) or 100 μM if not specified. LdtA, LdtB, and LdtC were expressed from pEP25, pAA186, and pEP26 respectively, without an inducer because no Lac repressor is present on the plasmids. However, when LppB was co-expressed with each Ldt from the two plasmids, 200 μM IPTG was added to guarantee the good expression of the Ldts. IPTG was added at the start of the culture. Cells were harvested at OD_600_ ∼ 0.5 by centrifuging at 9,400 x g.

### In vivo redox state determination of LppB

Cells were grown in LB medium until an OD_600_ of ∼ 0.5 and 1 mL of cells was precipitated by adding trichloroacetic acid (TCA) to a final concentration of 10% to preserve the thiol-disulfide redox states of proteins ^44^. When needed, samples were treated beforehand with 30 mM dithiothreitol (DTT) to reduce protein disulfide bonds and incubated on ice for 10 min before TCA precipitation. The TCA precipitated cells were washed with ice-cold acetone and solubilized in 100 mL SDS sample buffer (50 mM Tris-HCl, pH 8.0, 1% SDS, 10% glycerol, and 0.01% bromophenol blue) containing 10 mM N-ethyl maleimide (NEM, Sigma). NEM was used to alkylate free cysteines and prevents thiol-disulfide exchange reaction, allowing determination of in vivo thiol-disulfide redox states of proteins ^44^.

### Whole cell lysis and peptidoglycan digestion

Cells were grown in LB medium until an OD_600_ of ∼ 0.5 and 1 mL of cells were harvested. For experiments involving the expression of LppB, 10 mM N-ethylmaleimide (NEM) was added to the cell suspension to alkylate thiol groups and prevent thiol-disulfide exchange and cells were incubated on ice for 20 min. Cell were pelleted by centrifugation for 2 min at centrifuging at 9,400 x g and cell pellets were resuspended in 200 mL of lysis buffer containing 50 mM Tris-HCl (pH 8.0), 1 mM EDTA, 50 mM NaCl, 0.2% Triton X-100, and 1X protease inhibitor cocktail (Roche, catalog no. 11873580001). For peptidoglycan digestion, lysozyme (25 mg/mL; Sigma, catalog no. L1667) and mutanolysin (0.25 U/mL; Sigma, catalog no. M9901) were added to samples when necessary. Resuspended cells were incubated for 30 minutes at 30°C with shaking at 1500 rpm to facilitate cell lysis. Cells were subsequently precipitated with TCA, washed with ice-cold acetone and solubilized in 100 mL SDS sample buffer (see composition above) supplemented 30 mM DTT when necessary.

### Chemical crosslinking of Lpp or LppB and AMS alkylation

For experiments involving the expression of LppB, cells were treated with 1 mM of Dithiobis-succinimidyl-propionate (DSP, Covachem) or 2 mM of Bis-sulfosuccinimidyl-suberate (BS3, Covachem) and incubated at 30 °C for 30 min. After incubation, DSP and BS3 were quenched by adding glycine to a final concentration of 0.1M. Cells were subsequently precipitated with TCA, washed with ice-cold acetone and solubilized in 100 mL SDS sample buffer (see composition above) supplemented 30 mM DTT when necessary. The reduced BS3 samples were further alkylated by 4-Acetamido-4’-Maleimidylstilbene-2,2’-Disulfonic Acid (AMS, Invitrogen) ^44^.

### SDS-PAGE and Western blot analysis

Protein samples were separated via 10% or 12% SDS-PAGE (Life Technologies) and transferred onto nitrocellulose membranes (Amersham). The membranes were blocked overnight with 5% skim milk in TBS-T Buffer (50 mM Tris-HCl pH 7.6, 0.15 M NaCl, and 0.1% Tween 20). TBS-T Buffer was used in all subsequent immunoblotting steps. The primary antibody was diluted 1:200 in 1% skim milk in TBS-T Buffer and incubated with the membrane for 1h at room temperature. The membranes were incubated for 1 h at room temperature with horseradish peroxidase conjugated goat anti-rabbit IgG (Sigma) at a 1:10000 dilution in TBS-T Buffer. Antibody-labelled specific proteins were visualized by detecting chemiluminescence via the reaction of horseradish peroxidase (HRP) with luminol (Cytiva Amersham ECL Select).

### Antibody preparation

Polyclonal anti-Lpp antibodies were purified from rabbit antiserum (generated in the Collet laboratory with the CER Group, Marloie, Belgium) by affinity chromatography to the Lpp-derived peptide KVDQLSNDVNAMRSDVQAAK. 4,5 mg of synthetic peptide were conjugated in carbonate buffer (0.5 M NaHCO3 and 0.5 M NaCl, pH 8.3) to 320 mg of dry NHS-activated agarose resin (Pierce). 6 mL of serum containing polyclonal antibodies against Lpp were applied onto the affinity column and bound antibodies were eluted in in 50 mM glycine-HCl, 150 mM NaCl, pH 2.7. The eluate was immediately neutralized with 1 M Tris pH 7.5. About 180 μg of the purified antibody was obtained. The affinity-purified antibody was used as a primary antibody at a 1:200 dilution.

### Phylogenetic and genetic neighborhood analysis

Protein sequences were retrieved from the BioCyc Database Collections (EG10544 - G1G00- 1325 – BKC12_RS07715 – ECS_2384 - RS09425) and aligned using the MAFFT (v.7.511) program^45^. The resulting alignment was uploaded to the MEGA11 software to generate a maximum likelihood phylogeny tree ^46^. All genome sequences were downloaded from the NCBI Genome database (GCA_000005845.2 - GCA_000007545.1 - GCA_000006945.2 - GCA_000026545.1 -GCA_000008865.2) and uploaded to the Geneious Prime (v. 2024.0.7) software. The genomic regions of interest were extracted from the whole genome sequences and used to generate a genetic neighborhood comparison chart in Gene Graphics. The resulting diagram and the phylogeny tree chart were merged and refined using the BioRender software to enhance visual clarity and generate the final figure (Biorender.com).

### Sequence alignment

Protein sequences were retrieved from the BioCyc Database Collections (EG10544 - STM1377 - STM1376 - G7567) and aligned using the MAFFT (v.7) program ^45^. The 3D protein structure (X-Ray diffraction) of Lpp from *E. coli* was retrieved from PDB (1EQ7) and processed to generate the secondary structure depiction. The 3D protein structure (AlphaFold model) of YqhH from *E. coli* was retrieved from AlphaFold (AF-P65298-F1-v4) and processed to generate the secondary structure depiction. The resulting alignment was uploaded to the ESPript program (v.3.0) (https://espript.ibcp.fr/ESPript/ESPript/) to generate the alignment plot ^47^. The resulting chart was refined using the BioRender software to enhance visual clarity and generate the final figure (Biorender.com).

### SDS-EDTA sensitivity assay

Cells were grown in LB medium at 37 °C until they reached an OD_600_ of ∼ 0.5. Tenfold serial dilutions were made in LB medium and plated on LB agar pH 7 or pH 5.5, supplemented with 0.5 % SDS and 0.25 mM EDTA. Plates were incubated overnight at 37 °C.

### Structural modeling

The structure of the heterotrimer was predicted using AlphaFold3, as implemented on the AlphaFold Server available at https://alphafoldserver.com/. The resulting structure was manually duplicated and oriented using UCSF Chimera [https://pubmed.ncbi.nlm.nih.gov/15264254/], then the input system for the molecular dynamics simulation was prepared using the CHARMM-GUI webserver [https://pubmed.ncbi.nlm.nih.gov/18351591/] and the standard protocol for Membrane Builder/Bilayer Builder and CHARMM force field. The prepared system containing 157,424 atoms (including 35 LPS molecules for the outer leaflet and 60 PPPE, 16 PVPG and 4 PVCL2 lipids for the inner leaflet of the membrane) was equilibrated using NAMD [https://pubmed.ncbi.nlm.nih.gov/16222654/] with the standard CHARMM-GUI protocol for membrane proteins, then a production run was carried out for 100 ns. The simulation files were analyzed with MDAnalysis [https://pubmed.ncbi.nlm.nih.gov/21500218/] and the movie was generated using VMD [https://pubmed.ncbi.nlm.nih.gov/8744570/].

## ACKNOWLEDGEMENTS

We are grateful to K.T. Hughes (University of Utah) for providing us the Salmonella strains. We thank M. Deghelt for his helpful advice on the AMS alkylation experiment, D. Colau for antibody purification and F. García-del Portillo (CNB-CSIC, Madrid, Spain) for his helpful comments on the manuscript. This study was funded by the Fonds de la Recherche Scientifique FRS-FNRS (grant no. WELBIO-CR-2019C-03), the Fédération Wallonie-Bruxelles (grant no. ARC 22/27-128) and the Agence nationale de la recherche ANR (grant no. ANR-20-PAMR-0010).

## AUTHOR CONTRIBUTIONS

E.S.P.D., S.-H.C. and J.-F.C. wrote the paper; S.-H.C., and J.-F.C. conceived the project, E.S.P.D., S.-H.C. and J.-F.C. designed the study; E.S.P.D., S.-H.C., B.I.I. performed the experiments; E.S.P.D., S.-H.C., B.I.I. and J.-F.C. analyzed the data.

## DATA AND MATERIALS AVAILABILITY

All data are available in the main text or the supplementary materials or upon request. Correspondence and requests for materials should be addressed to Jean-François Collet.

## SUPPLEMENTARY FIGURES

**Figure S1.**
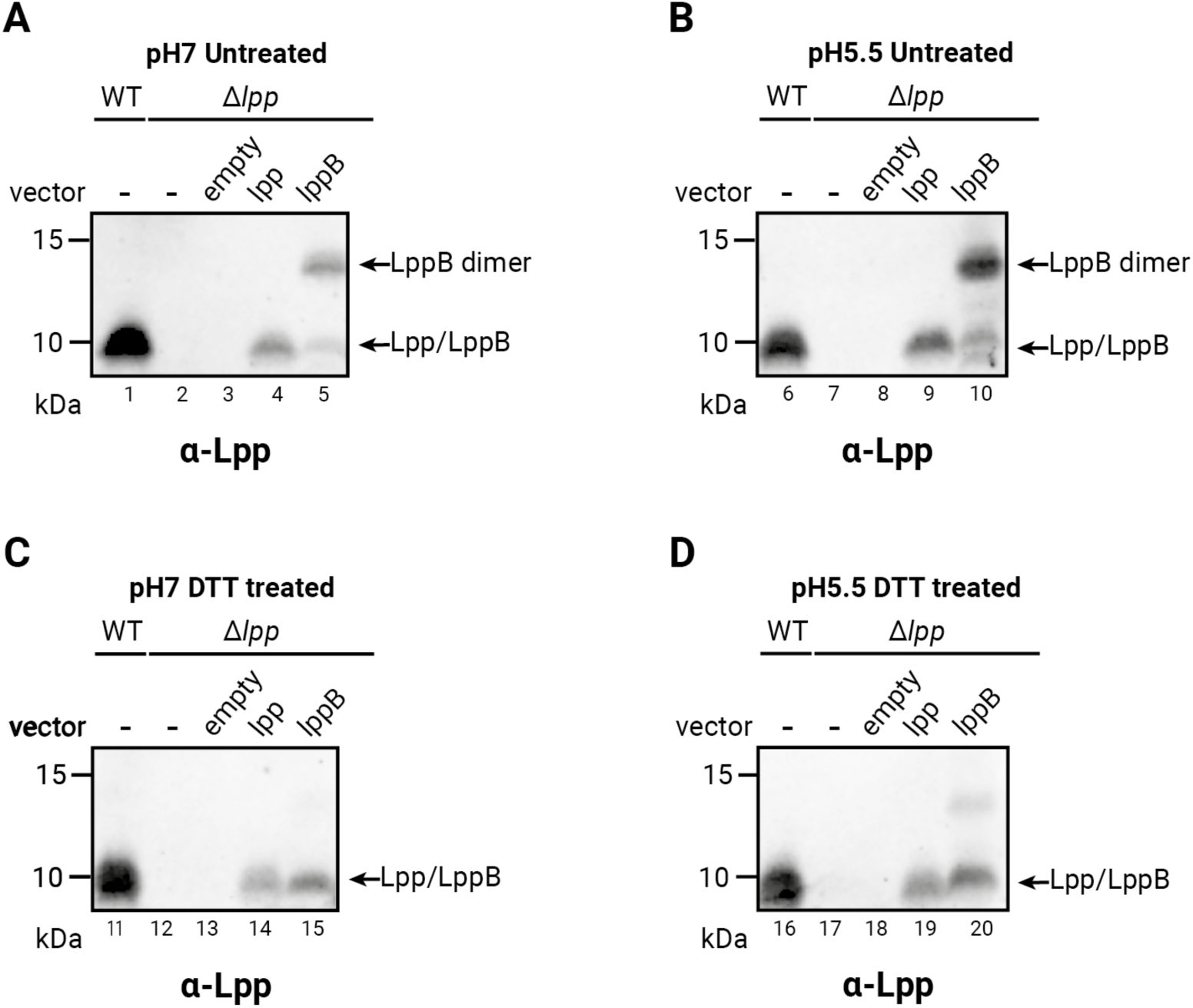
Acidic pH increases LppB expression levels. Lpp and LppB variants were expressed from pSC284 and pSC283 derivatives respectively, in the Δ*lpp* strain. **A.** Lpp and LppB oligomeric states in neutral pH conditions. Untreated cell lysates from cells grown at pH 7 show monomeric Lpp and LppB at ∼7.5 kDa and LppB homodimers at ∼15 kDa. **B.** Lpp and LppB oligomeric states in acidic pH conditions. Untreated cell lysate from cells grown at pH 5.5 reveal higher LppB expression compared with neutral conditions (compare lanes 5 and 10). Monomeric and dimeric species migrate as in panel A. **C. - D**. Effect of reducing conditions. Lysates from panels A and B were treated with 30 mM DTT to reduce disulfide bonds. DTT treatment decreased LppB dimer abundance and increased monomer signal at both pH conditions, confirming disulfide bond–dependent dimerization. Western blot analysis was performed using anti-Lpp antibody. Representative images from experiments conducted in biological triplicates are shown.

**Figure S2.**
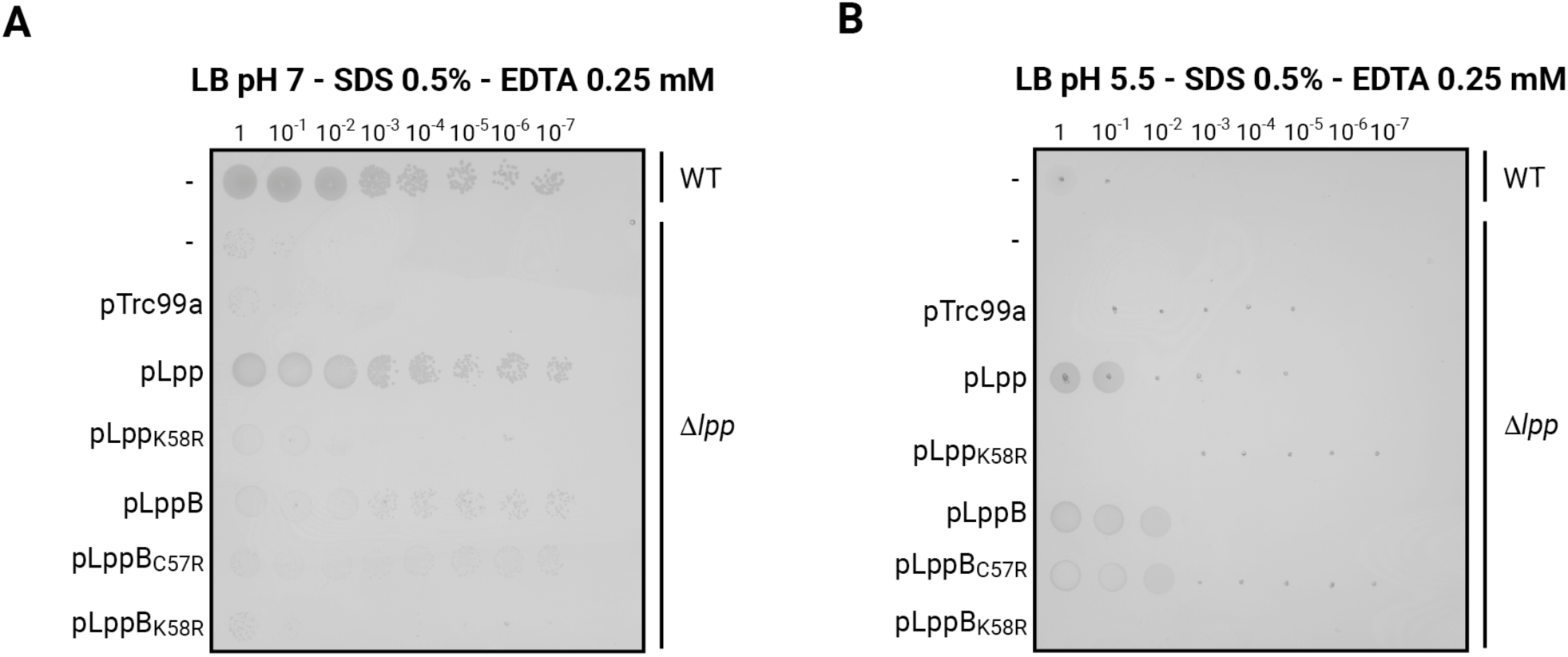
Acidic pH enhances LppB-mediated protection against SDS-EDTA sensitivity. The contribution of LppB to envelope integrity increases under acidic conditions. Wild-type and Δ*lpp* mutant complemented with pSC284 and pSC283 derivatives were grown in LB at the indicated pH. Cells were harvested, serially diluted, and 5 μL of each dilution was spotted on indicated plates and incubated overnight at 37 °C. **A.** At pH 7, LppB expression provides weak protection against SDS-EDTA-induced growth defects compared to Lpp. **B.** At pH 5.5, LppB expression restores resistance to SDS-EDTA-induced growth defects to levels comparable to those conferred by Lpp. Representative images from experiments conducted in biological triplicates are shown.

**Figure S3.**
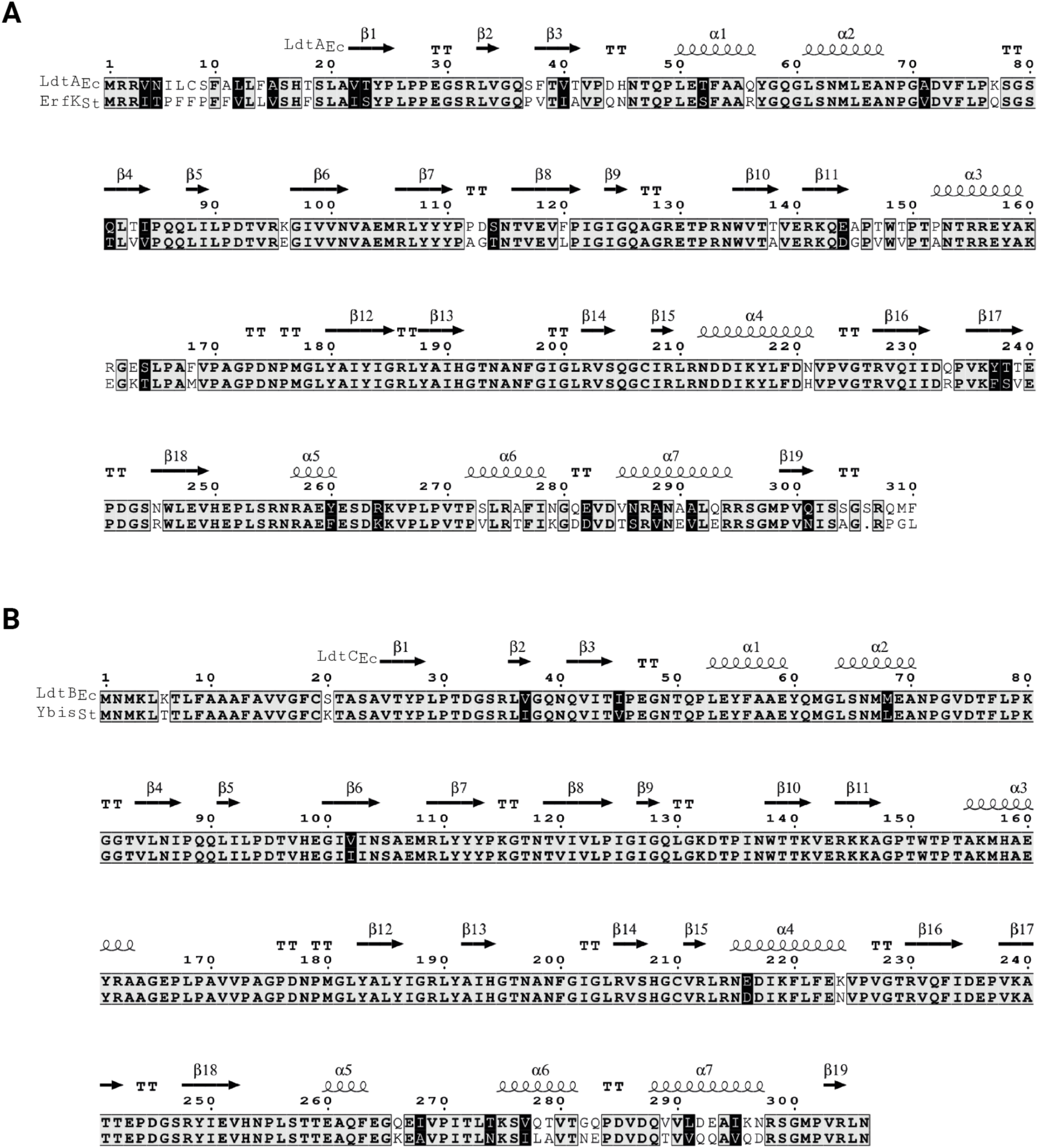
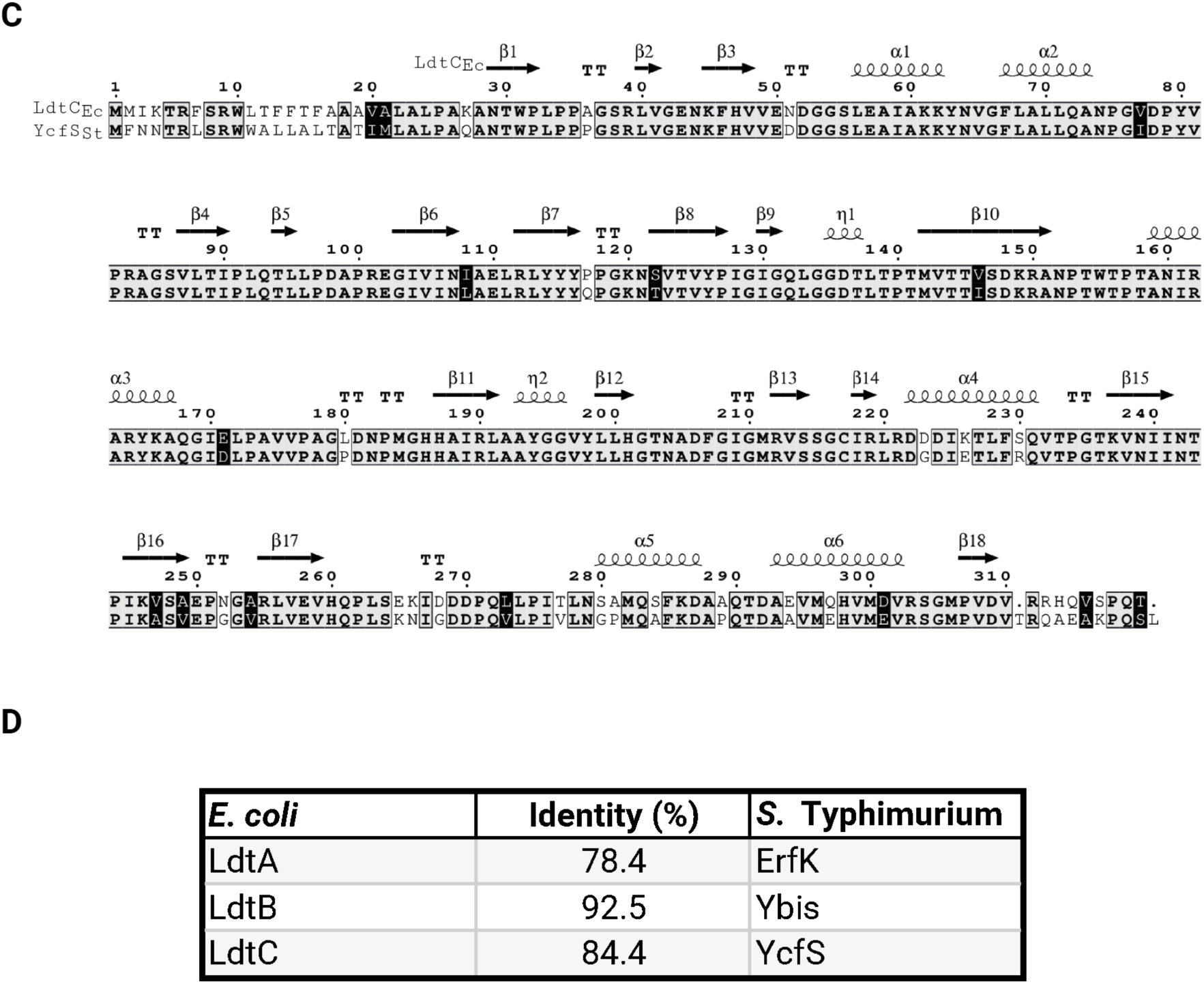
*E. coli* LdtA/B/C share strong sequence similarity with their *S.* Typhimurium homologs ErfK/Ybis/YcfS. Amino acid sequence alignments of the L,D-transpeptidases from *E. coli* K12 and *S.* Typhimurium LT2. Numbers above the first sequence line indicate amino acid positions relative to the start of each protein sequence. Conserved residues are shown as black letters on a grey background, and similar residues are shown as white letters in black boxes. **A.** Amino acids sequence alignment of *E. coli* LdtA and *S.* Typhimurium ErfK. The secondary structure of LdtA is depicted above the alignment. **B.** Amino acids sequence alignment of *E. coli* LdtB and *S.* Typhimurium YbiS. The secondary structure of LdtB is depicted above the alignment. **C.** Amino acids sequence alignment of *E. coli* LdtC and *S.* Typhimurium YcfS. The secondary structure of LdtA is depicted above the alignment. **D.** Pairwise percent identity matrix. Percent identity values were calculated from global pairwise alignments performed with EMBOSS Needle and represent the proportion of identical residues across the full alignment length, including gaps.

**Figure S4.**
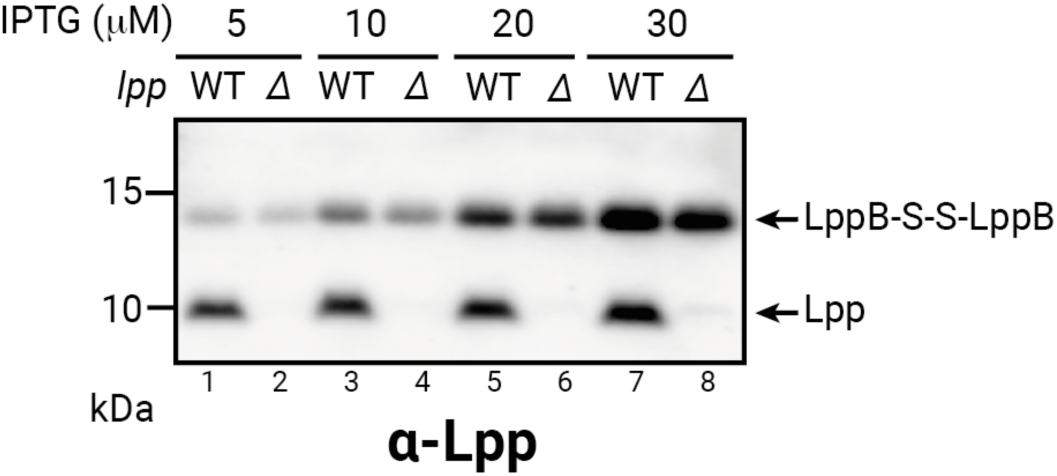
LppB forms homodimeric mixed disulfide independently of expression levels or Lpp presence. LppB was expressed from pSC283 (pLppB) in the wild-type and *Δlpp* strains using varying concentrations of isopropyl ß-D-1-thiogalactopyranoside (IPTG) for induction. Bands corresponding to the homodimeric mixed-disulfide form of LppB (∼15 kDa) were detected under all tested conditions indicating that LppB homodimerization occurs independently of its expression level or the presence of Lpp. Western blot analysis was performed using anti-Lpp antibody. Representative images from experiments conducted in biological triplicates are shown.

**Figure S5.**
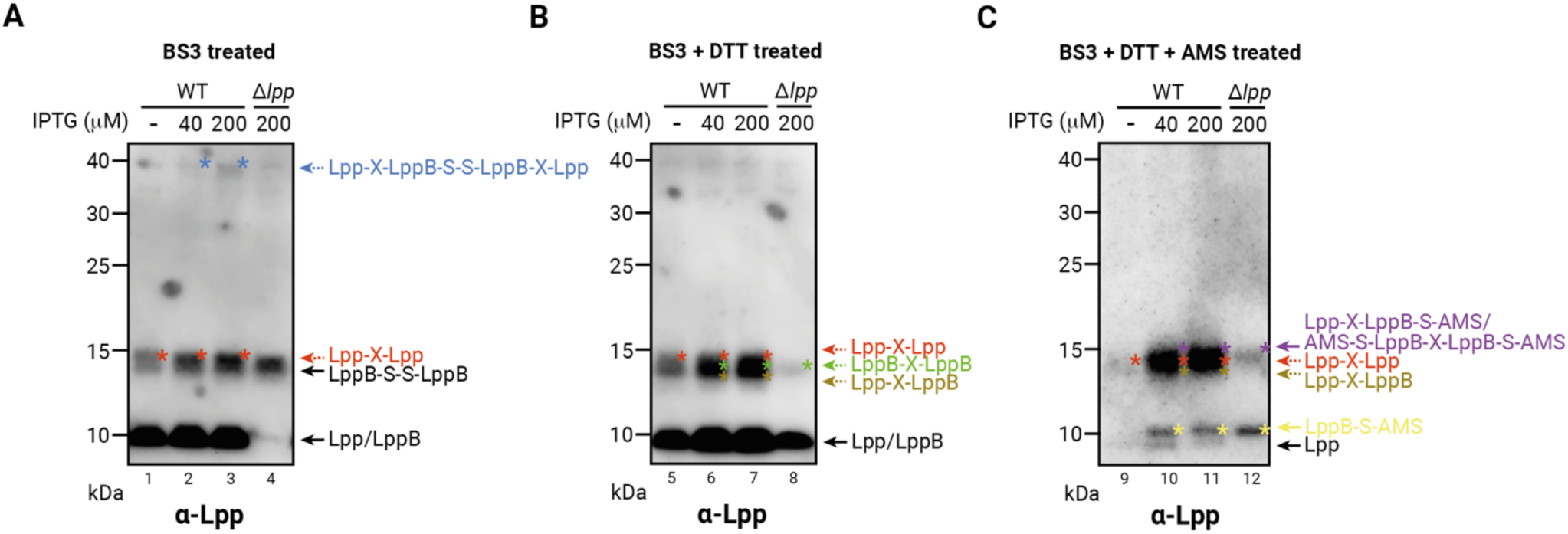
LppB forms homodimeric mixed disulfide and heterotrimers with Lpp in vivo. LppB was expressed from pSC283 (pLppB) in the wild-type and *Δlpp* strains using varying concentrations of isopropyl ß-D-1-thiogalactopyranoside (IPTG) for induction. **A.** Chemical crosslinking with Bis-sulfosuccinimidyl-suberate (BS3) reveals distinct oligomeric states of Lpp and LppB. Bands corresponding to Lpp or LppB monomers and homodimers and to Lpp-LppB heterotetramers (Lpp-LppB-S-S-LppB-Lpp) were detected. The samples analyzed here are identical to those used in Fig. 5C. **B.** Effect of reducing conditions on BS3-crosslinked Lpp and LppB oligomeric states. Samples from panel A were treated with 30 mM DTT to reduce disulfide bonds formed through native cysteine only. Disulfide bond formed through BS3 are irreversible. The samples analyzed here are identical to those used in Fig. 5F. **C.** Effect of alkylating conditions on reduced BS3-crosslinked Lpp and LppB oligomeric states. Reduced samples from panel B were treated with 4-Acetamido-4’-Maleimidylstilbene-2,2’-Disulfonic Acid (AMS), which adds ∼ 500 Da per free thiol. AMS treatment caused mobility shifts for LppB dimers and Lpp-LppB heterodimers, but not for Lpp homodimers, confirming the presence of reactive cysteines in LppB. Western blot analysis was performed using anti-Lpp antibody.

**Figure S6.**
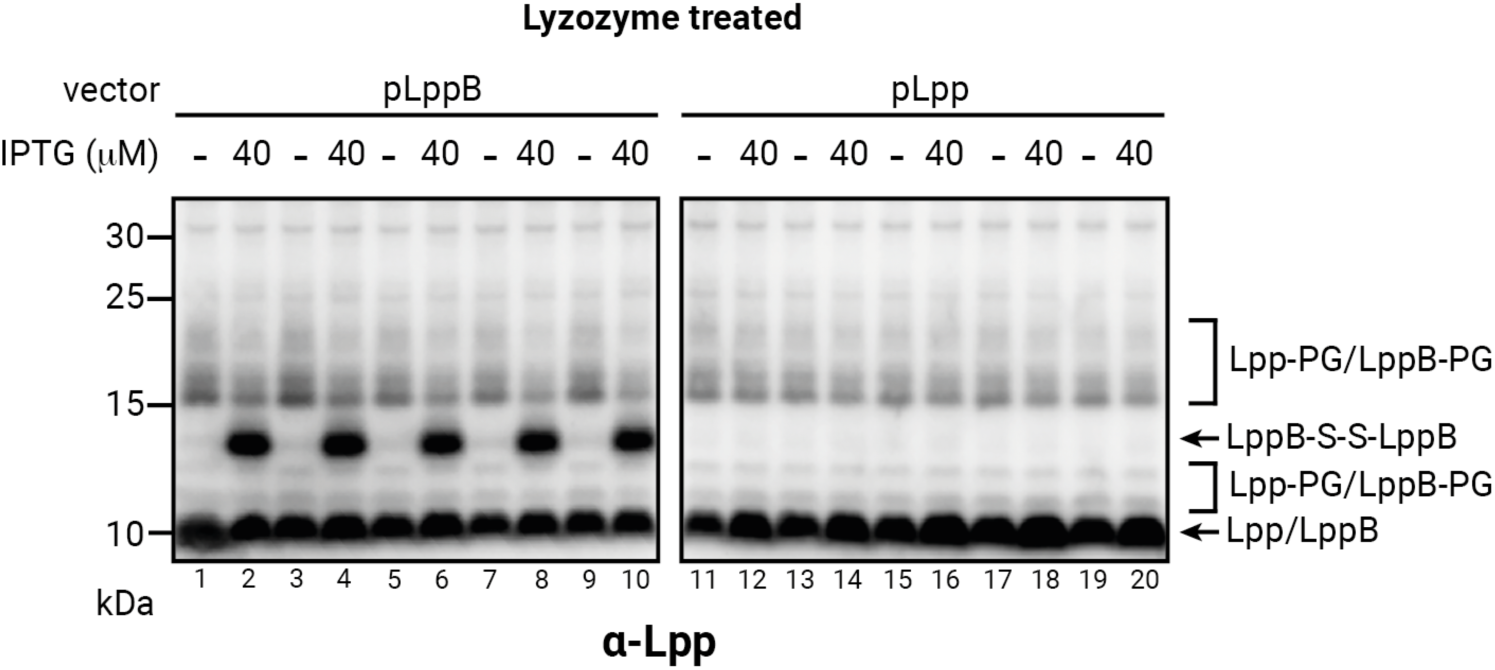
LppB negatively modulates Lpp-peptidoglycan crosslinking. Comparative effects of LppB and Lpp on Lpp-peptidoglycan crosslinking. LppB was expressed from pSC283 (pLppB) and Lpp from pSC284 (pLpp) in wild-type cells using the indicated concentrations of isopropyl β-D-1-thiogalactopyranoside (IPTG) for induction. The left panel shows LppB expressed at 0 or 40 μM IPTG; the right panel shows Lpp expressed under the same conditions. Increasing LppB expression reduced Lpp-peptidoglycan crosslinking, whereas Lpp overexpression had no detectable effect. Cells were harvested and treated with lysozyme for PG digestion. Western blot analysis was performed using anti-Lpp antibody, and representative images from experiments conducted in biological triplicates are shown.

**Figure S7.**
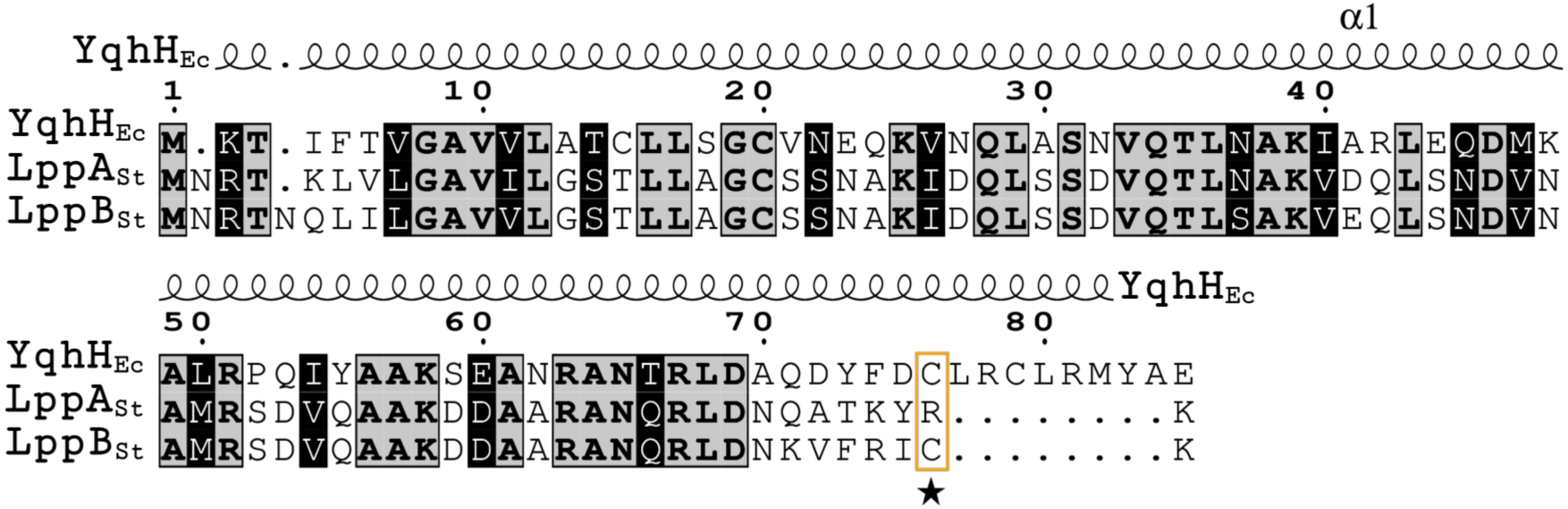
E. coli Lpp paralog YqhH shares sequence similarity with S. Typhimurium LppA and LppB, and a conserved C-terminal cysteine is found in YqhH and LppB. Amino acid sequence alignment of *E. coli* YqhH with LppA and LppB from *S.* Typhimurium LT2. YqhH contains a conserved cysteine residue at its C-terminus that aligns with Cys57 of LppB (highlighted within an orange box and marked by a star). The predicted secondary structure of *E. coli* YqhH is depicted above the alignment. Illustration created with ESPript and Biorender.com.

## SUPPLEMENTARY DATA

### Statistical modeling of LppA-LppB trimer formation

LppA and LppB share strong sequence homology in their coiled-coiled helices, whereas the C-terminal region exhibits sequence variations outside the coiled-coil domain (Fig. 1B and Fig. 6H). This suggests that LppA and LppB can form heterotrimers, as proteins are unlikely to discriminate each other when folded. This hypothesis is supported by the results obtained by chemical crosslinking experiments (Fig. 5B-F; Fig. S5).

Based on this, we modeled the proportional distribution-of LppA and LppB heterotrimers mathematically. In principle, the abundance of the four possible trimers —two homotrimers (AAA, BBB) and two heterotrimers (AAB, ABB)— depend on the relative concentrations of both LppA (A) and LppB (B) in the cell.

To explore the role of LppB as a limiting factor, and since LppB is expected to be present at low levels in Salmonella, we calculated trimer probabilities as a function of the LppB fraction (b), with the fraction of LppA equal to 1 – b. Under the assumption that trimers form *without preference* for homotrimers, the probabilities of forming each trimer would be:

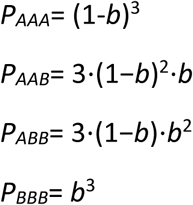

Since LppB is expected to be present at low levels *in vivo*, P_AAA would be predicted to be the largest, followed by P_AAB, whereas P_ABB and P_BBB would be negligible. For example, if b = 0.1, then P_AAA = 0.729, P_AAB = 0.243, P_ABB = 0.027, and P_BBB = 0.001.

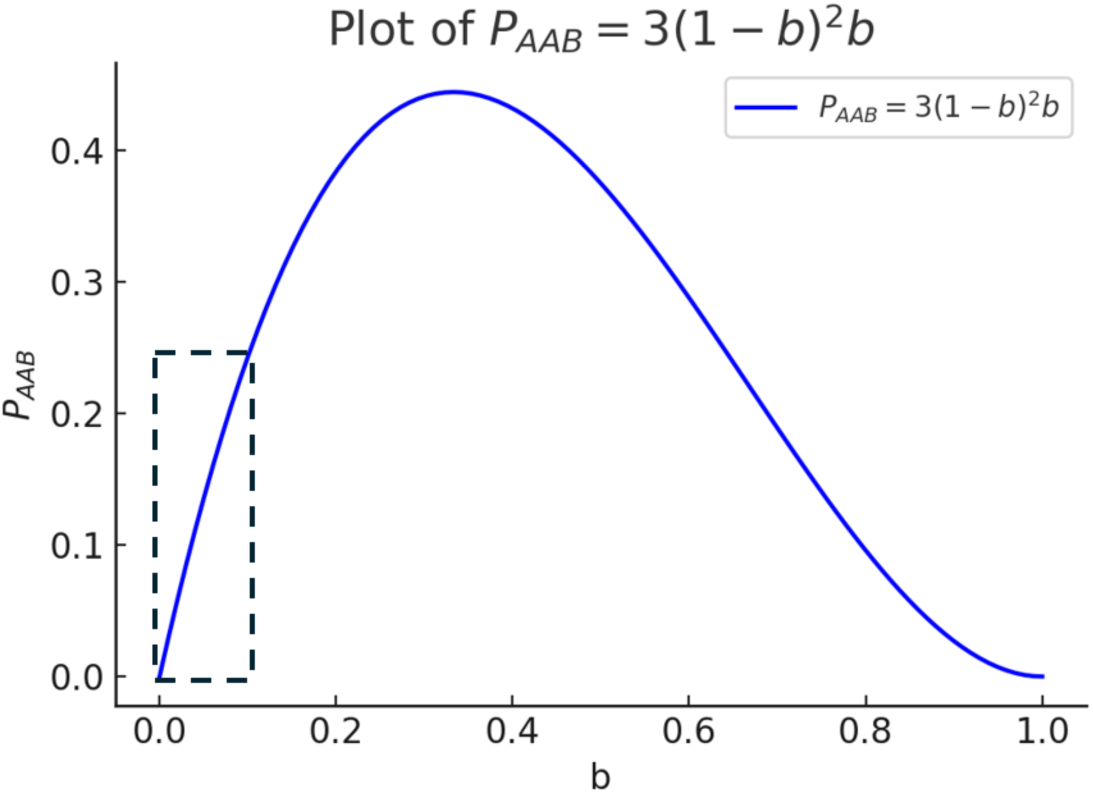
Formation of LppA and LppB heterotrimers. The plot shows the probability of AAB heterotrimer formation (*P_AAB)_* as a function of the fraction of LppB (*b)*. *P_AAB_* increases up ∼ b=0.33, then declines as the relative fraction of LppA decreases. Since LppB is expected to be present at low levels in Salmonella, the region where b is small (highlighted by the dotted square) is particularly informative. Within this range, *P_AAB_* increases approximately linearly, with a ∼ 2.4-3-fold rise as the fraction of LppB increases

## SUPPLEMENTARY MOVIES

**Supplementary Movies.**
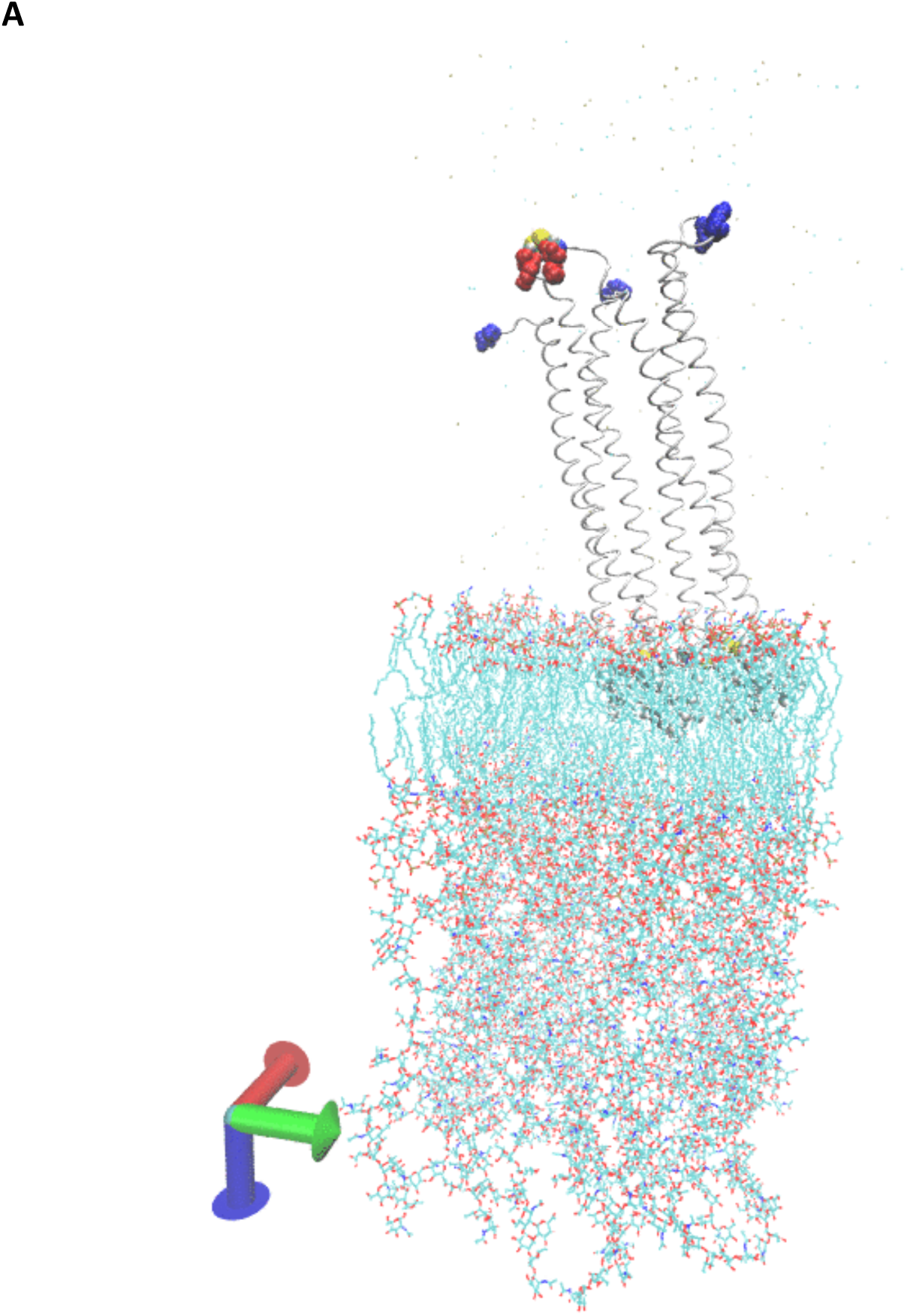
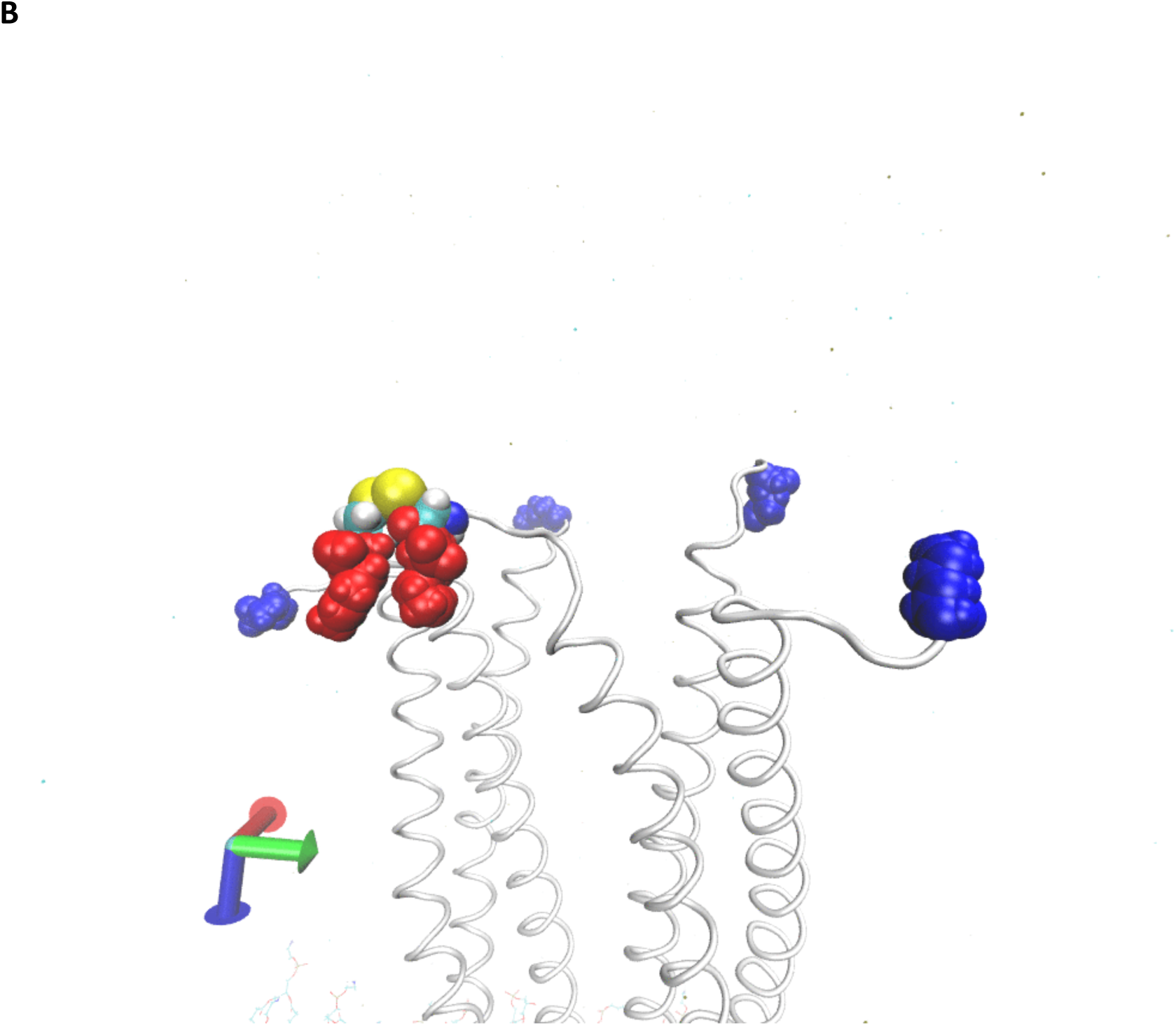
Molecular dynamics simulation depicting a homodimeric mixed disulfide complex comprising two LppA copies and one LppB copy [LppA2LppB]-S-S-[LppBLppA2] anchored in the outer membrane (OM). **A.** Terminal lysine and cysteine residues are represented as spheres (sulfur atoms in yellow; Lpp Lys58 in blue; LppB Lys58 in red). Lpp/LppB helices appear as ribbons, with N-terminal acyl chains embedded in the OM (shown as sticks). **B.** A magnified view of (A) focusing on the Cys58 residues in LppB and the Lys58 residues in both Lpp and LppB.

## SUPPLEMENTARY METHODS

### Sequence alignment and identity percentages calculations

Protein sequences were retrieved from the BioCyc Database Collections (G7073 - G6422 - G6571 - STM2015 - STM0837 - STM1215) and aligned using the MAFFT (v.7) program ^1^. The 3D protein structures (AlphaFold model) of LdtA, LdtB and LdtC from *E. coli* were retrieved from AlphaFold (AF-P39176-F1-model_v4, AF-P0AAX8-F1-model_v4, AF-P75954-F1-model_v4) and processed to generate the secondary structure depiction. The resulting alignment was uploaded to the ESPript program (v.3.0) (https://espript.ibcp.fr/ESPript/ESPript/) to generate the alignment plot ^2^. The resulting chart was refined using the BioRender software to enhance visual clarity and generate the final figure (Biorender.com). Pairwise sequence alignment was performed using EMBOSS Needle with the EBLOSUM62 substitution matrix, a gap-open penalty of 10.0, and a gap-extension penalty of 0.5 ^3^. For the LdtA_Ec_/ErfK_St_ pair, the resulting global alignment between the two proteins was 310 amino acids in length and showed 78.4% identity (243/310) and 86.8% similarity (269/310), with one gap (0.3%). The alignment score was 1305.0. For the LdtB_Ec_/Ybis_St_ pair, the resulting global alignment between the two proteins was 306 amino acids in length and showed 92.5% identity (283/306) and 96.7% similarity (296/306), without gap (0.0%). The alignment score was 1511.0. For the LdtC_Ec_/YcfS_St_ pair, the resulting global alignment between the two proteins was 321 amino acids in length and showed 84.4% identity (271/321) and 89.7% similarity (288/321), with one gap (0.3%). The alignment score was 1421.0.

### SDS-EDTA sensitivity assay

Cells were grown in LB medium at 37°C until they reached an OD_600_ of ∼ 0.5. Tenfold serial dilutions were made in LB medium and plated on LB agar pH 7 or pH 5.5, supplemented with 0.5 % SDS and 0.25 mM EDTA. Plates were incubated overnight at 37 °C.

### Methodology for Plotting *P_AAB_*

To visualize the function *P_AAB_*=3(1−*b*)^2^*b*, we used the following computational approach. Firstly, the mathematical expression *P_AAB_* =3(1−*b*)^2^*b* was implemented as a function in Python ^4^ . Second, the variable *b* was sampled within the range [0,1] using a uniform grid of 100 points.

Third, the plot was generated using Python with the NumPy library for numerical computation and Matplotlib for visualization ^5,6^. Fourth, **t**he function *P_AAB_* was evaluated at each point in the specified range. A line plot was created with *b* on the x-axis and *P_AAB_* on the y-axis. Labels, a title, and a grid were added to improve readability. Fifth, the computations were performed in a Python environment using Python version 3.x, libraries of NumPy (for array computations) and Matplotlib (for plotting), and resolution. The function was evaluated at 100 equally spaced points in the interval [0,1]

## SUPPLEMENTARY TABLES

**Table S1.**
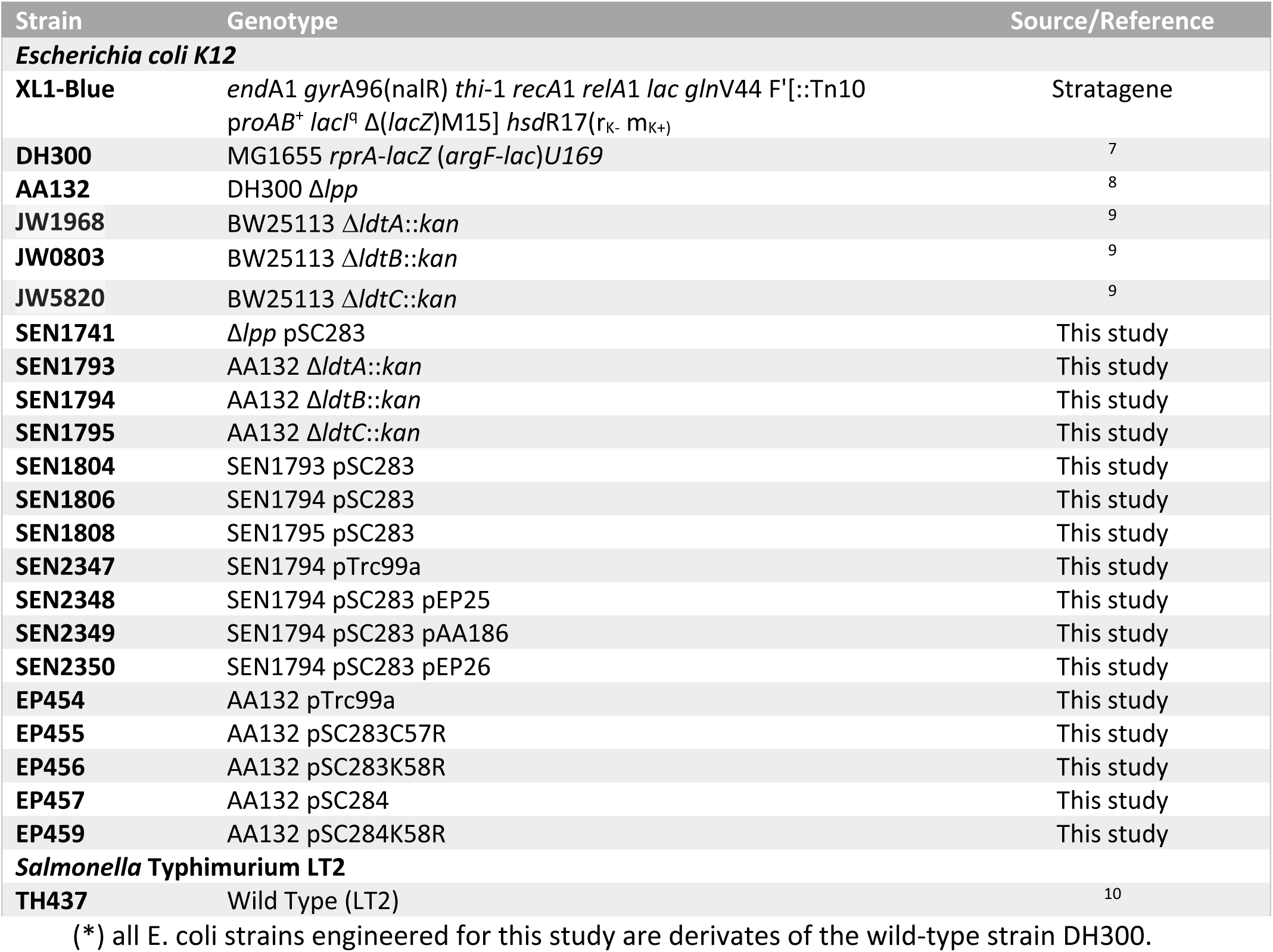
Bacterial strains used in this study.

**Table S2.** Plasmids used in this study.

| Plasmid | Features | Source/Reference |
| --- | --- | --- |
| **pAM238** | IPTG-regulated $P_{lac}$, pSC101-based, spectinomycin<sup>R</sup> | 11 |
| **pTrc99a** | IPTG-regulated $P_{trc}$, pBR322-origin, ampicillin<sup>R</sup> | Pharmacia |
| **pCP20** | $FLP^+$, $\lambda$ cl857<sup>+</sup>, $\lambda_{PR}$ Rep<sup>ts</sup>, ampicillin<sup>R</sup>, chloramphenicol<sup>R</sup> | 12 |
| **pSC283** | pTrc99a_StLppB | This study |
| **pSC284** | pTrc99a_EcLpp | This study |
| **pSC284K58R** | pTrc99a_EcLpp Lys58Arg | This study |
| **pSC283C57R** | pTrc99a_StLppB Cys57Arg | This study |
| **pSC283K58R** | pTrc99a_StLppB Lys58Arg | This study |
| **pEP25** | pAM238_LdtA | This study |
| **pAA186 (pYbiS)** | pAM238_LdtB | 8 |
| **pEP26** | pAM238_LdtC | This study |

(\*)antibiotic<sup>R</sup> indicates that the vector carries a resistance marker for the specified antibiotic.

**Table S3.**
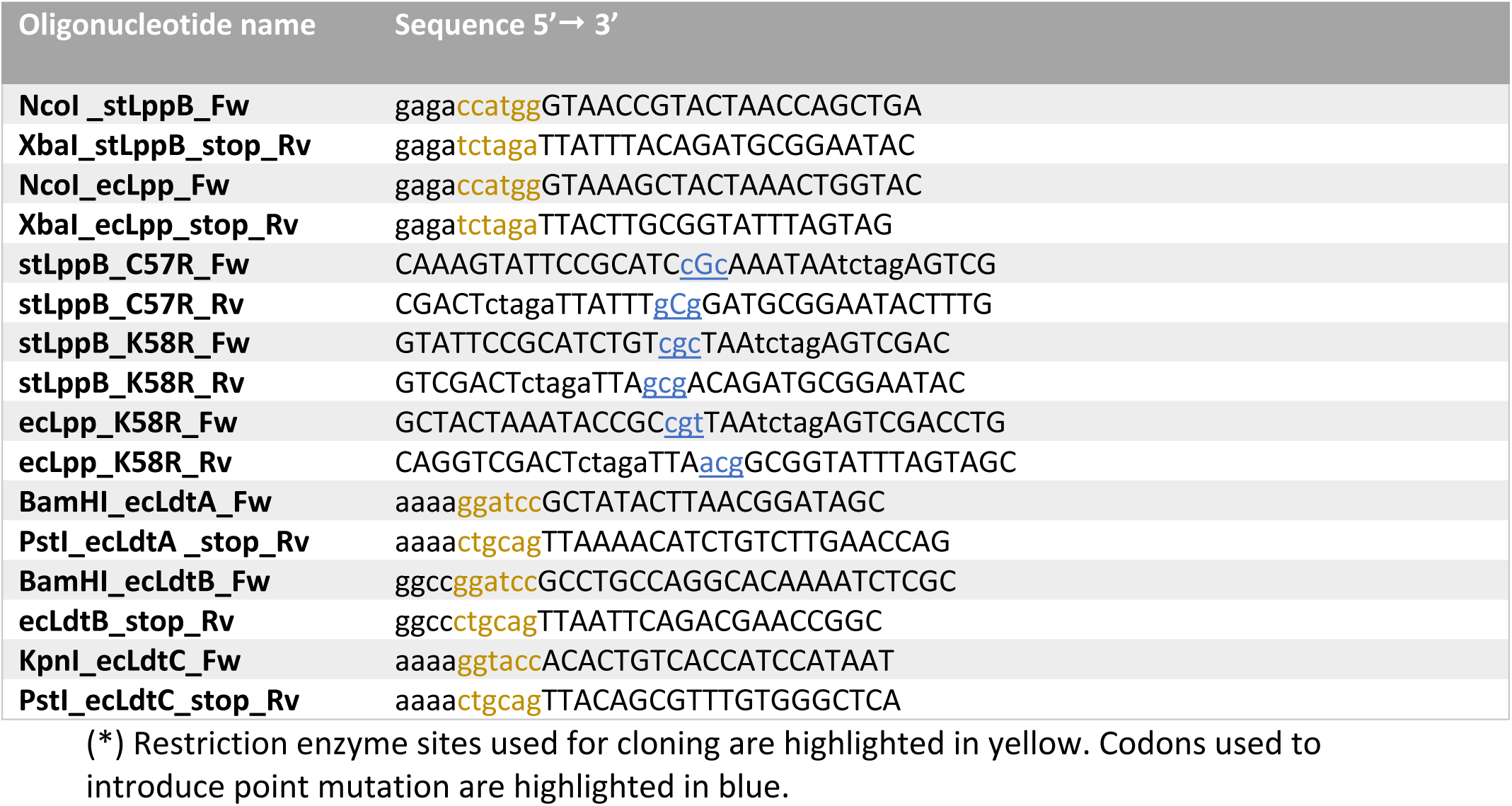
Primers used in this study.

